# A chemical perturbation scan on a K_2P_ channel modulator pocket identifies affinity hotspots

**DOI:** 10.64898/2026.10.13.758835

**Authors:** Edward Mendez-Otalvaro, Bert L. de Groot

## Abstract

Ion permeation in the two-pore-domain K^+^ (K_2P_) channel TREK1 is gated largely at the selectivity filter (SF), by a C-type-like inactivation mechanism involving distortions of the SF structure; and by a carbonyl flip, which disrupts K^+^ coordination. Both are prevented by a small-molecule activator bound adjacent to the SF, in a site termed the modulator pocket; yet which pocket regions carry the binding signal, and whether they couple to gating, is unclear.

We use 46 chemical substitutions on the activator ML335, evaluated by non-equilibrium alchemical free-energy calculations, as an affinity perturbation scan on the lower-and upper-pocket. The response is strongly asymmetric: Lower-ring substitutions impair binding, which we trace to a *π*-*π* stacking network connecting the M3, P2, P1 helices, and the SF. Conversely, upper-ring substitutions are tolerated and reshape contacts with the Cap–P1 and SF2–M4 loops; and with the upper SF. The identified affinity hotspots overlap with residues independently implicated in TREK gating by external cues such as pH, temperature, pressure, and cytosolic-sensor transduction, suggesting that high-affinity pocket sites might also shape the gating landscape.

## Introduction

TWIK-related potassium channel 1 (TREK1, IUPHAR Nomenclature K_2P_2.1) is a two-pore domain potassium (K_2P_) channel that plays a critical role in regulating the resting membrane potential in human nervous tissue.^3,4^ Due to its function, TREK1 is getting significant therapeutic interest for the treatment of nerve disorders associated with cellular hyperexcitability and pain transmission.^5–8^

Structurally, TREK1 is a dimer composed of four transmembrane helices (M1-M4), two pore-forming helices (P1 and P2), and a selectivity filter (SF), that is, the narrowest region of the pore that determines ion selectivity. The SF consists of two strands, SF1 and SF2, which display slight amino acid variations, resulting in an asymmetric filter within each subunit.^3,9^

Experimental and computational studies indicate that TREK1 gating primarily occurs at the SF. The channel alternates between an open state, with the SF adopting a canonical conductive conformation, and a C-type-like inactivated state, in which the SF shows a distorted geometry that hinders ion flow.^10–12^ In addition, it has been proposed that TREK1 permeation can be altered by the flipping of a carbonyl oxygen that forms the third K^+^ binding site (S3) on SF1 (a conformational change that introduces a significant energetic barrier to ion permeation).^11^

More recently, Lolicato *et al.*^9^ reported two activators, ML335 (Q6F) and ML402 (Q5F), that enhance the activity of TREK1. These activators have EC_50_ values of 10.5 *±* 2.7 *µ*M (Q6F) and 14.9 *±* 1.6 *µ*M (Q5F) in functional assays measuring current intensity in the crystal constructs. Structurally, both ligands bind at the interface between the top of the M4 helix and the P1 helix in a cryptic site termed the “modulator pocket” by the authors. This pocket is located adjacent to the SF1 strand, with each subunit of the dimer accommodating one ligand molecule^9^ (**Figure S1**).

Interestingly, previous evidence suggests that specific hydrophobic and polar motifs regulate the activation of TREK1 and other K_2P_ channels via the modulator pocket. ^12^ For example: (i) binding of the anionic lipid 1-palmitoyl-2-oleoyl-sn-glycero-3-phosphatidic acid (POPA) to the entrance of this pocket activates TREK1;^13^ (ii) a fatty acid molecule, bound within a sub-region of the equivalent pocket, acts as a structural component of the conductive state in THIK1;^14^ (iii) a nanobody (Nb-Activator-76) induces conformational changes in TREK2 that stabilize the side chains of residues W275 (M4 helix) and F134 (P1 helix); which are homologous to key modulatory residues in TREK1, promoting channel activation; ^15^ and (iv) molecular modelling, mutagenesis, and electrophysiological studies show that the negatively charged molecule C3001a binds to a region overlapping the TREK1 modulator pocket, resulting in channel activation.^16^

The mechanism by which Q6F activates TREK1 involves a reduction in SF distortions, which stabilizes its canonical conductive conformation and prevents C-type-like inactivation.^10^ This stabilization might be mediated by interactions between residues N147 and F134, located at the top of SF1 and P1, respectively.^11^ In addition, the presence of Q6F appears to counteract the S3 carbonyl-flip mechanism of inactivation by stabilizing the side chain of a residue located at the base of SF1 (T141), which in turn reinforces the backbone conformation of the S3 and reduces the local water occupancy within and behind this binding site.^11^

Furthermore, Q6F selectively modulates TREK over other K_2P_ channels, such as TRAAK,^9^ through a critical *π*–cation interaction mediated by its upper phenyl ring. Removing this interaction, either by aliphatic substitution of the phenyl ring (Q6Fa) or by neutralizing the corresponding cationic residue in TREK1 to mimic TRAAK, abolishes binding. However, despite its selectivity, Q6F affinity for TREK1 (reflected in its micromolar EC_50_ range) suggests that there is room for improvement in potency. Given that C-type inactivation and carbonyl flipping are mechanisms that may be modulated by ligand binding, ^12^ such improvements could be tailored to these gating processes.

Therefore, in light of previous experimental evidence and the two proposed gating mechanisms in TREK1 (C-type-like inactivation and carbonyl flipping) we hypothesize that introducing more hydrophobic and sterically larger groups on the Q6F lower phenyl-ring could enhance binding affinity and potentially modulate the S3 flipping mechanism. Likewise, incorporating hydrophilic or charged groups on the Q6F upper phenyl-ring may improve binding while also helping to prevent C-type inactivation of the channel. This hypothesis is motivated by the amphipathic character of the modulator pocket: hydrophobic features in the lower region and hydrophilic features in the upper region of such pocket appear to be a conserved physicochemical and structural motif across K_2P_ channels,^12^ and complementing this conserved architecture from the ligand perspective is an opportunity to improve binding, establishing in that way affinity hotspots in the pocket that, besides, might be coupled to gating (**Figures 1 & S2**).

**Figure 1:**
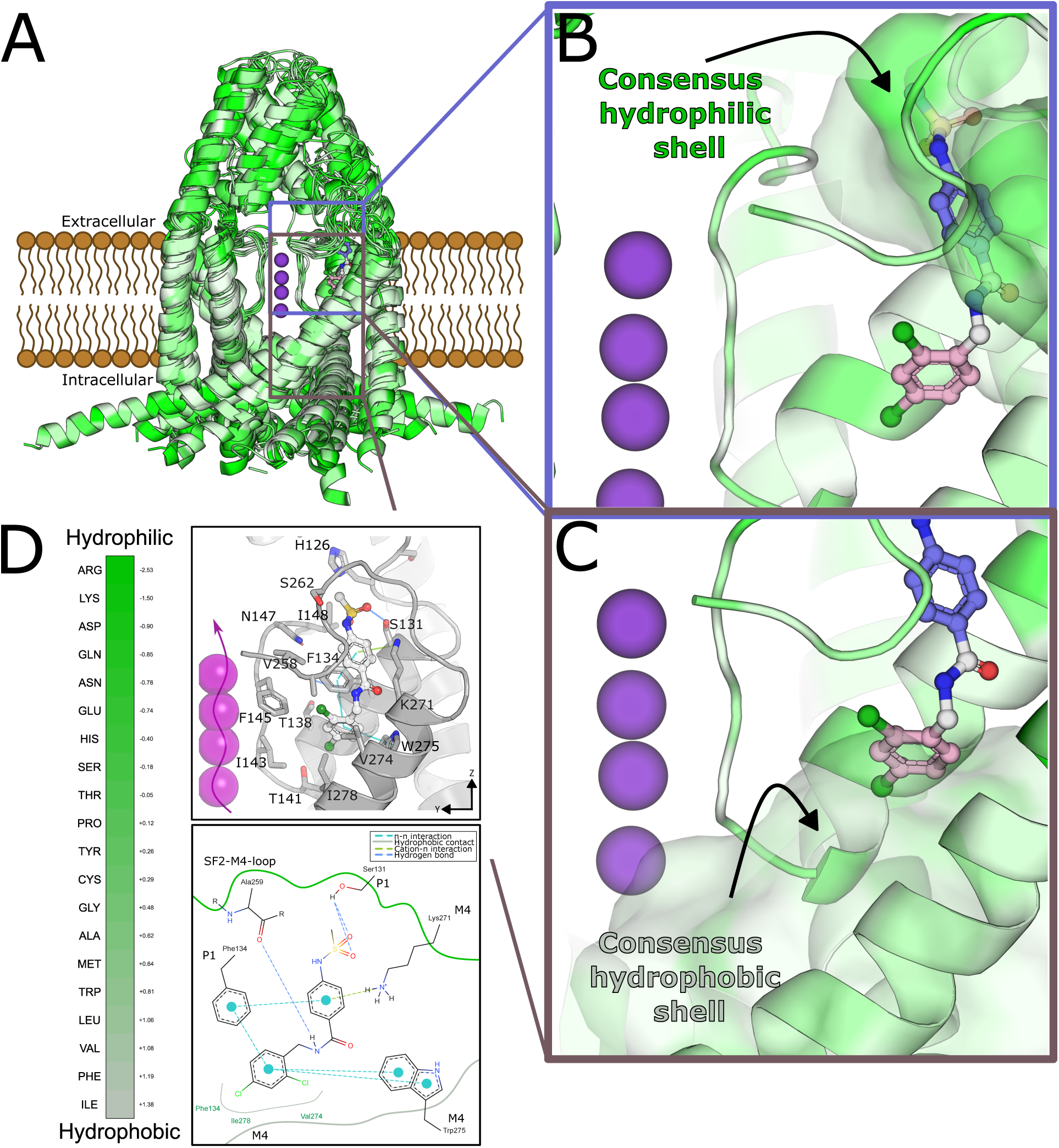
Hypothesized ligand and channel features contributing to affinity in the TREK1 modulator pocket. **Panel A**. Structural superposition of experimentally resolved K_2P_ channel structures aligned around the transmembrane region and SF. Q6F is shown as sticks in the modulator pocket, and K^+^ ions in the SF as purple spheres. Blue and dark-pink boxes indicate the upper and lower pocket regions analyzed in Panels B and C. **Panel B**. Upper pocket region, with the upper aromatic ring of Q6F shown in blue and the transparent surface representing the consensus hydrophilic environment across the aligned K_2P_ structures. **Panel C**. Lower pocket region, with the lower aromatic ring of Q6F shown in pink and the transparent surface representing the consensus hydrophobic environment. **Panel D**. Eisenberg hydrophobicity scale ^1^ used to color the aligned structures and surfaces. Lower sub-panel: Two-dimensional representation ^2^ of the main non-covalent interactions of Q6F (chain A) in the modulator pocket, based on the TREK1 holo x-ray structure (PDB: 6CQ8). Upper sub-panel: Zoomed *zy*-plane view of the Q6F binding pose in the PDB model embedded in a solvated membrane simulation box, with key residues from the P1 helix, M4 helix, and SF1 region labeled.

To probe how the pocket responds to those changes, we treat a chemical series as a perturbation scan. Because the perturbation is applied to the ligand rather than to the protein, a region in which substitution is strongly penalized is then one that is saturated in the bound state, whereas a tolerant region retains spare capacity available for optimal binding. In this way, we computed relative binding free energies for 46 Q6F derivatives using rigorous non-equilibrium statistical mechanics, calibrated against experimental and structural data for Q6F, Q5F and the non-binder Q6Fa, and analyzed the contact patterns of the derivatives with the channel across the equilibrium ensembles.

Our results indicate that affinity hotspots within the pocket are asymmetric: upper-ring perturbations are generally better tolerated, whereas lower-ring substitutions are more detrimental to binding. This suggests that affinity depends on more than the modulator pocket itself: Favorable binding is associated with contacts involving the SF2-M4 loop, M3-P2 interface, and upper SF1 region, while the M4 region and central pocket act as hydrophobic anchors whose disruption is detrimental. Further, these regions overlap with previously reported gating residues in TREK channels.

These trends are reflected in the chemistry of the substitutions: Moderate lipophilicity, distributed polarizability, and mildly electron-rich surfaces favor binding, whereas bulky groups, localized heavy atoms, and electropositive lower-ring surfaces tend to destabilize binding and reduce affinity.

## Methodology

### Computational models

Structures of the TREK1 channel in the apo state (PDB: 6CQ6), and in holo states bound to ML335 (PDB: 6CQ8) or ML402 (PDB: 6CQ9) were obtained from the Protein Data Bank (PDB).^9^ Unresolved loops in chain A were modeled using the respective resolved regions in chain B and vice versa. All systems were protonated at pH 7.0, capped with acetyl (ACE) and N-methyl amide (NME) groups, and embedded in a 1-palmitoyl-2-oleoyl-snglycero-3-phosphocholine (POPC) membrane, surrounded by water, chlorine (Cl^-^) and K^+^ ions, forming a hexagonal box with a salt concentration of approximately 0.5 M using the CHARMM-GUI web server,^17^ which provides a pre-equilibrated initial membrane system.

For the free energy calculations, solvated ligand systems were constructed separately in rectangular simulations boxes containing explicit water and Cl^-^/K^+^ ions at the same nominal concentration (0.5 M), using GROMACS 2024.4.^18^ The box dimensions were chosen to ensure a minimum distance of at least 2.0 nm between the solute and the box boundaries.

### Simulation and free energy calculations setup

Simulations were performed using GROMACS 2024.4.^18^ Proteins, lipids, water, and ions were modeled using the CHARMM36m,^19^ CHARMM36,^20,21^ CHARMM TIP3P,^22^ and Beglov & Roux parameters,^23^ respectively. All small molecules were parameterized using CGenFF^24–26^ obtained from CHARMM-GUI. For experimentally validated molecules (Q6F, Q5F, and Q6Fa), any parameters with a penalty score greater than 50 were refined using the ffTK as implemented in VMD^27^ (**Figures S3-S6**). CHARMM force fields were obtained directly from the developers’ distribution (charmm36-jul2021.ff), while CgenFF parameters were generated using CHARMM-GUI.^28^

Since the Gibbs free energy is a thermodynamic state function, only the free energy difference between two states A and B matters, not the path connecting them. This allows the construction of thermodynamic cycles in which the total free energy change is zero. Accordingly, instead of directly computing the binding free energy difference between a reference ligand and a derivative (which is computationally demanding), we compute the free energy of transforming the reference ligand into the derivative, both in solution and in the bound state. This alchemical approach is computationally more tractable and estimates the same target relative binding free energy difference.

Therefore, the thermodynamic cycle consisted of two branches (protein and water) and two states (reference and modified) (**Figure S7**). In the protein branch, the reference state corresponds to the Q6F ligand conformation in the modulator pocket of TREK1 chain A, while the modified state displays the Q6F derivative at the same site. The modulator pocket in chain B remained unmodified, retaining the native Q6F ligand conformation. In the water branch, a ligand with hybrid topology (Q6F or derivative, depending on the alchemical state) was simulated in a fully solvated water box with the same ionic strength as in the protein branch (**Figure S7**).

In order to computationally implement the two Hamiltonians (Q6F ligand and derivative) in the simulation; the single topology approach was used for all free energy calculations.^29^ Ligand topologies and hybrid structures (Q6F ligand and derivative) were generated using the *pmx* tool,^30,31^ specifically the ligandHybrid and atomMapping commands. The resulting hybrid structure was inserted into the pre-equilibrated box provided by CHARMM-GUI.

For each state in the thermodynamic cycle, five independent replicates were carried out using the following protocol: i.) Energy minimization (steepest descent algorithm with maximum 10 000 steps). ii.) NVT equilibration for 10 ns with position restraints on protein, ligand, and phospholipid atoms (*k_pr_^xyz^* = 1000 *kJ · mol^−^*^1^ *· nm^−^*^2^). iii.) NPT equilibration for 20 ns with position restraints on protein and ligand (*k_pr_^xyz^* = 1000 *kJ ·mol^−^*^1^ *·nm^−^*^2^). iv.) Production run in the NPT ensemble for 50 ns without restraints. v.) Non-equilibrium transitions: The first 26 ns of the production run were discarded as equilibration. From the remaining 24 ns, 300 frames were extracted. For each frame, fast transitions of 1 000 ps were performed in both directions, where the system’s Hamiltonian is coupled to a *λ* parameter that interpolates the ligand non-physically from the reference to the modified state and vice versa. This results in 600 transitions per replicate. During these transitions, soft-core potentials were applied to both Lennard-Jones and Coulomb interactions using the Gapsys method, in order to avoid singularities due to overlapping atoms during the decoupling-coupling phase. ^32^

All simulations employed the leapfrog stochastic dynamics integrator^33^ with a 2 fs time step. Pressure was maintained at 1 bar using the Berendsen barostat^34^ for equilibration and C-rescale barostat^35^ for production (semi-isotropic or isotropic, depending on the branch), and the temperature was kept at 298.15 K using Langevin dynamics ^36^ (as included in the sd integrator). Electrostatic interactions were computed using Particle Mesh Ewald (PME)^37^ with a real-space cutoff of 1.2 nm, spline order 4, relative tolerance of 10^-5^, and Fourier grid spacing of 1.2 nm. Van der Waals interactions were force-switched off between 1.0 and 1.2 nm, and h-bonds were constrained using the LINCS algorithm.^38^

Some ligands required modifications to this protocol: For the experimentally validated ligands (Q6F, Q5F, and Q6Fa), which displayed larger structural perturbations, the transition time was increased to 2 000 ps to improve convergence. For charged ligands, artifacts can arise when computing electrostatics using PME. ^39^ To address this, a co-alchemical counterion was introduced, interpolated simultaneously with the ligand to maintain box neutrality. This ion was position-restrained near the box edge, akin to the double-system/single-box setup.^31^

Free energy differences were calculated by integrating the derivative of the Hamiltonian with respect to the *λ* parameter, yielding the dissipated work along each transition.^31^ Forward and backward work distributions were then used to estimate free energy differences via a maximum likelihood estimator based on the Crooks fluctuation theorem^40,41^ (**Figure S7**), as implemented in the pmx analyse tool.^31^ Uncertainties were reported as bootstrap-estimated standard errors of the mean across the five independent replicates.

### Chemical library construction and pose generation

The fragment library was constructed from Enamine’s Hit Locator Library (460 160 compounds, representing a commercially available HTS-ready collection) as follows (**Figure S8**): The N-benzoylbenzylamine scaffold from the Q6F activator was used to identify ligands with this topological substructure within the Enamine library using RDKit. ^42^ Molecules displaying the substructure were then listed and compared to Q6F by calculating graph-based RDKit fingerprints and computing their Tanimoto similarity. This process produces 1 059 ligands from the initial library that were both highly similar to Q6F and contained the N-benzoylbenzylamine scaffold.

Next, the crystallographic pose of the Q6F ligand bound to TREK1 chain A was used to generate three-dimensional coordinates of these 1 059 molecules using RDKit’s ConstrainedEmbed algorithm.^42^ Briefly, the N-benzoylbenzylamine scaffold served as the substructure match between Q6F and each ligand. The ligand was then minimized using the UFF force field while constraining the scaffold atoms to match the coordinates of the Q6F conformation. This resulted in a minimized structure with a conformer constrained to the binding pose.

Given the diversity of substitutions on both the upper (N-phenylmethanesulfonamide) and lower (1,3-dichlorobenzene) rings (and their potential overlap with residues in the modulator pocket) we generated two sub-libraries containing only the fragments attached to each respective ring. Substitutions that overlapped with any atom in the modulator pocket (based on the crystallographic structure) were discarded. This filtering produced two fragment libraries of 269 and 224 fragments for the upper and lower rings, respectively, forming the chemical space to be explored for Q6F substitutions (**Figure S8 Panel A**). Molecules that might fit after slight rearrangements or induced fit effects are not included in this approach.

The initial poses of the Q6F derivatives were prepared as follows (**Figure S8 Panel B**): Fragments selected to test our hypotheses were mapped onto the 3D structures generated in the previous step. The aligned system was subjected to energy minimization (maximum 10 000 steps) with position restraints (*k_pr_^xyz^* = 1000 *kJ · mol^−^*^1^ *· nm^−^*^2^) applied to all heavy atoms of the protein and the N-benzoylbenzylamine scaffold, allowing the derivative’s ring substitutions to move freely.

The minimized structure was then processed using a simulated annealing protocol, where the temperature was decreased from 298.15 K to 0 K over 1 000 ps, with the same position restraints applied. The final pose from this “freezing” protocol was used as the initial structure for subsequent simulations. The minimization and simulated annealing protocol were carried out using GROMACS 2024.4

### Order parameters measured from the equilibrium ensembles

A set of order parameters was computed from the equilibrium ensembles sampled for each derivative in order to identify those degrees of freedom that correlate with improved binding affinity. These descriptors are summarized as follows:

- **Coordination number (CN)**: Defined as the number of water oxygen atoms within a distance of 3.5 Å of any ligand heavy atom. This quantity is computed both in the solvent ensemble (w) and in the channel-bound ensemble (p).
- **Total contact fraction, hydrophobic contact fraction (HCF), and polar contact fraction (PCF)**: A contact is defined when any atom of a given selection group lies within a distance of 4.5 Å of any ligand heavy atom. The selection groups considered are: (i) total contacts, defined as all heavy atoms of the protein; (ii) hydrophobic atoms, defined as carbon atoms belonging to residues ALA, VAL, ILE, LEU, MET, PHE, TYR, TRP, and PRO; and (iii) polar atoms, defined as nitrogen, oxygen, sulfur, or phosphorus atoms belonging to residues SER, THR, ASN, GLN, GLY, ARG, HIS, LYS, ASP, GLU, TYR, and TRP. The hydrophobic and polar contact counts are normalized by the total number of contacts per frame to obtain HCF and PCF, respectively.
- **Conformational entropy**: For all derivatives, five dihedral angles around the amide bond linking the upper and lower phenyl rings in Q6F were measured: *φ*_1_ : *C*_1_ *− C*_2_ *− N*_1_ *− H*_13_, *φ*_2_ : *O*_1_ *− C*_2_ *− N*_1_ *− H*_13_, *φ*_3_ : *C*_10_ *− C*_8_ *− N*_1_ *− H*_13_, *φ*_4_ : *H*_9_ *− C*_8_ *− N*_1_ *− H*_13_, and *φ*_5_ : *H*_8_ *− C*_8_ *− N*_1_ *− H*_13_. These dihedrals describe the conformation of the N-methylacetamide group, a motif conserved across all derivatives. The probability distributions of these dihedral angles are computed, and the Gibbs entropy (*S* = *−k_B_∑_i_ p_i_* ln *p_i_*) is evaluated for each distribution. Summing these contributions one obtains an estimate of the torsional entropy associated with these degrees of freedom for each ligand. The difference between the bound and solvated states provides a proxy for the conformational entropy penalty upon binding.

For all order parameters, a bootstrapped standard error of the mean (SEM) was estimated across five independent replicates.

### Total contact fraction analysis

To understand how the interaction patterns of the derivatives differ from that of Q6F, we first computed, for each derivative, the average total contact fraction with each channel residue across replicates, as explained in the previous section. Since each derivative displays a different interaction profile across residues, we then summarized the contact matrix by taking, for each ligand-residue pair, the maximum fraction of total contacts observed across the ligand heavy atoms. Finally, this matrix was expressed relative to the corresponding matrix of the reference ligand Q6F. In this way, for each derivative, we get an interaction-pattern matrix that quantifies the change in maximum residue contact fraction with respect to Q6F.

## Results

### A chemical perturbation scan on the modulator pocket reveals an asymmetric tolerance to substitution in terms of affinity

Before testing or refuting our hypotheses regarding the amphipathic nature of the modulator pocket as a determinant of activator affinity for TREK1, we first estimate relative binding free-energy differences corresponding to experimentally characterized cases (**Figures 2 & 3, orange and green colors**). Specifically, we considered: (i) the relative affinity difference between Q6F and Q5F, which serves as a positive binding control;^9^ and (ii) the relative affinity difference between Q6F and Q6Fa, which works as a negative binding control, as Q6Fa does not exhibit measurable affinity for TREK1 due to the removal of the upper aromatic ring.^9^ From these comparisons, we observe that our computational estimates fall within the same order of magnitude as the experimentally derived affinity proxies. Although these experimental values (reported as EC_50_ measurements) are themselves only approximations to the true standard binding free energy, our approach successfully recapitulates the expected qualitative trends: distinguishing between a binding and a non-binding molecule. Together, these benchmarks indicate confidence that our computational estimates probe a physically meaningful range of relative affinities, and support their use for ranking the Q6F derivatives explored in this work.

**Figure 2:**
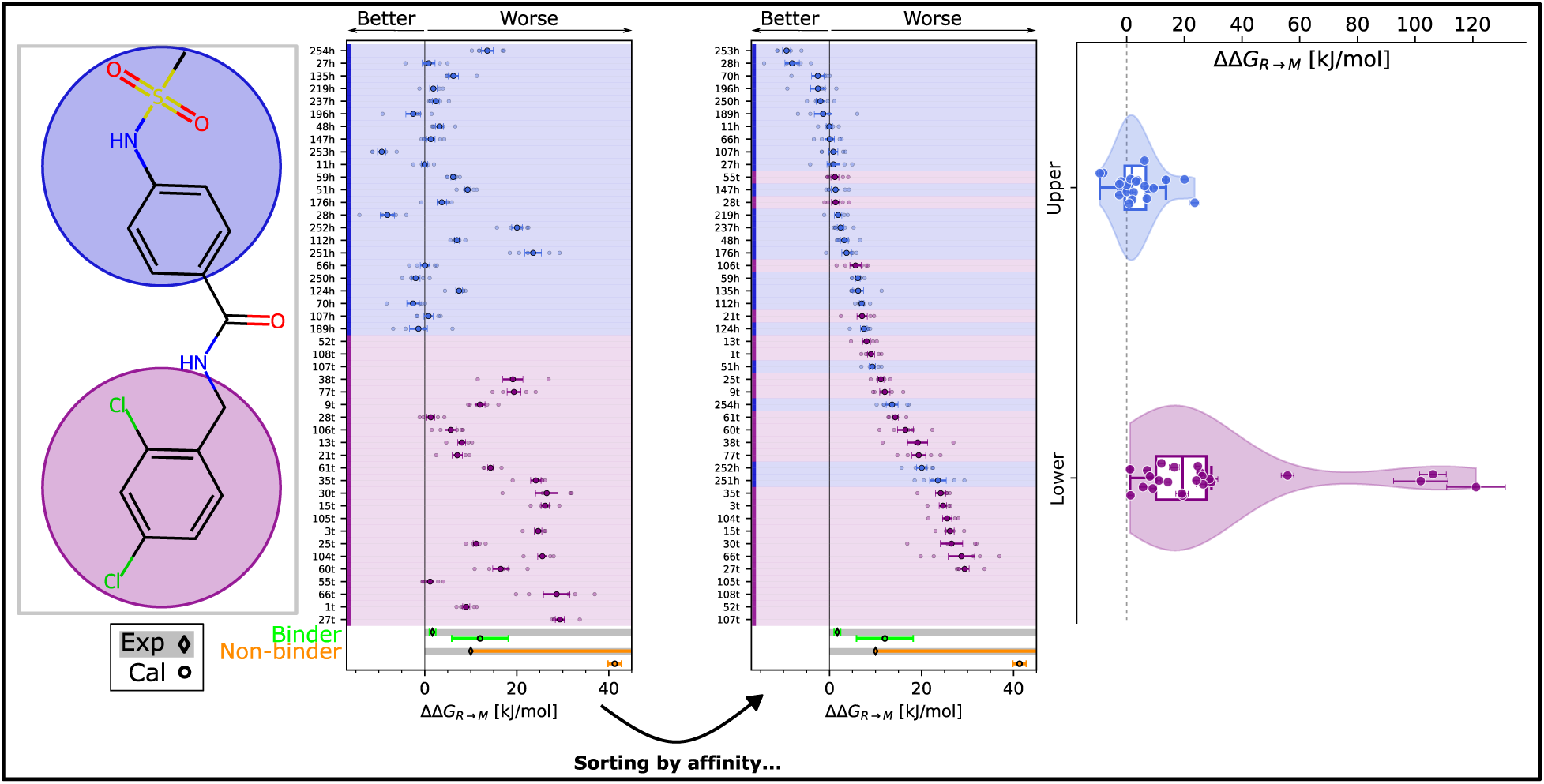
Relative binding free energy differences reveal the impact of substitutions in Q6F. **Left panel:** Structure of the Q6F molecule, with the upper and lower phenyl rings highlighted in pastel blue and light pink, respectively. The background colors indicate the substituted ring. **First middle panel**: Relative binding free energy differences across all chemical perturbations; the corresponding derivatives are shown in Figure 3. Error bars represent the bootstrapped standard error of the mean, and the individual points correspond to replicate calculations. The labels *t* and *h* denote substitutions at the tail (lower ring) and head (upper ring) of the molecule, respectively. The relative binding free energy differences for a binder (Q6F *→* Q5F) and a non-binder (Q6F *→* Q6Fa) are shown in green and orange, respectively. Both experimental (diamond markers) and computationally calculated values (circle markers) are displayed. The experimental value for the non-binder does not have a well-defined error estimate, as no saturation was observed in the dose–response curve. ^9^ Consequently, the molecule does not exhibit measurable biological activity, and its experimental EC_50_ is not well defined. **Second Middle panel**: Same as in the middle panel, but with the relative binding free energy differences sorted by their mean values. More negative values indicate stronger binding relative to the reference (Q6F). **Right Panel**: Distributions of relative binding free energy calculations for the total perturbation dataset, discriminated based on upper- or lower-ring substitutions.

**Figure 3:**
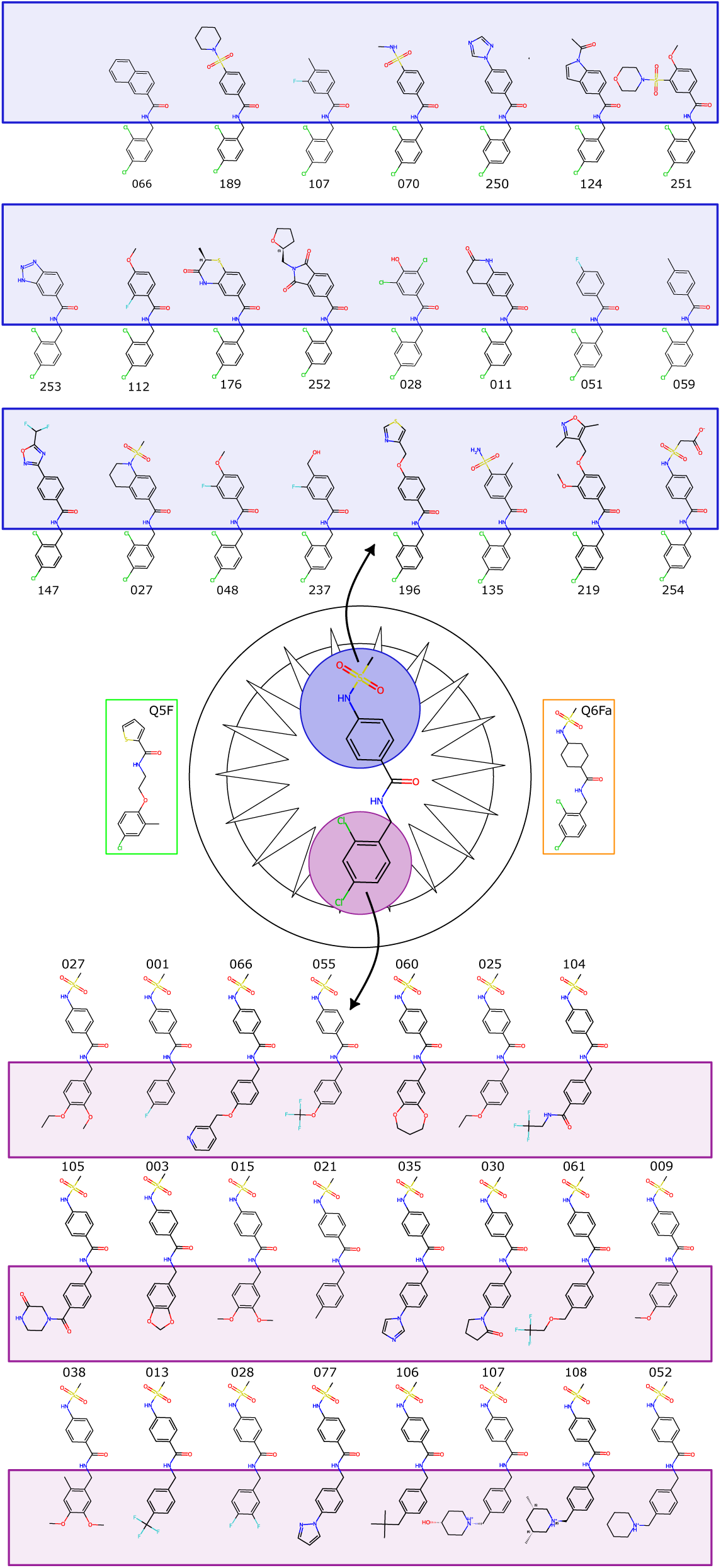
Derivatives used for the chemical perturbation. Q6F derivatives with substitutions on the upper ring (pastel blue) and lower ring (light pink) are shown. The numbering is consistent with Figure 2. The chemical space sampled includes derivatives spanning a range of sizes, hydrophobicity, charge, degree of ramification, steric hindrance, polarity, and related physicochemical properties. The experimentally validated binder (Q5F), and non-binder (Q6Fa) are highlighted in green and orange squares, respectively.

In our iterations of systematic modifications to the upper and lower aromatic rings of Q6F, we observed an unexpected trend: substitutions on the upper ring perturb the relative affinity of Q6F for TREK1 to a lesser extent than substitutions on the lower ring (**Figures 2 & 3**). Our initial hypothesis proposed that the lower pocket represents a more conserved hydrophobic motif among K_2P_ channels, such that substitutions increasing the interaction surface with residues lining this region might enhance affinity. However, our results suggest that modifications at this position instead might disrupt a relatively well-preserved interaction motif, in which the Q6F-phenyl ring engages residues F145, F244, and F134 through a *π*–*π* stacking network that is already well optimized. We will expand further about this interesting result later.

Modifications to the lower ring fail, on average, to improve the relative affinity of Q6F for TREK1. In contrast, substitutions on the upper ring more effectively engage interaction hotspots within the upper modulator pocket, enabling additional stabilizing contacts and, consequently, improving binding affinity. Collectively, these results highlight the asymmetric contribution of the two aromatic rings to ligand binding, and identify the upper ring as a more optimal and promising site for affinity optimization (**Figures 2 & 3, Table S1**). Still, we noted that some lower-ring substitutions yielded large positive ΔΔG values (*∼*5 SD above the group mean), so we examine the convergence of each transformation as a function of sampling time (**Figures S1A-S5A**) and the overlap of the forward and reverse non-equilibrium work distributions (**Figures S6A-S10A**). For most derivatives both legs are converged and the work distributions overlap, supporting the quantitative use of the corresponding estimates.

For the three most unfavorable lower-ring compounds (107t, 108t and 52t), however, the protein-leg work distributions have low overlap, so the Crooks estimator is poorly determined and the associated values (*>*100 kJ mol*^−^*^1^) should not be read as calibrated affinities. We instead interpret the poor work overlap as a model-independent signal that these substituents cannot be accommodated in the lower pocket without a large reorganization of the bound state (which is the qualitative conclusion we draw from them throughout).

### Upper-ring substitutions preserve a more consistent protein-ligand contact pattern and are associated with improved binding affinity compared to the lower-ring ones

So far, we have observed an asymmetric affinity response to chemical perturbations of the Q6F activator. To get more insight about the origin of this result, we analyzed ligand-pocket contacts by measuring, for each molecule, the fraction of the maximum number of contacts with the channel pocket. Differences relative to Q6F reveal how chemical modification reshapes the contact profile (**Figure 4**).

**Figure 4:**
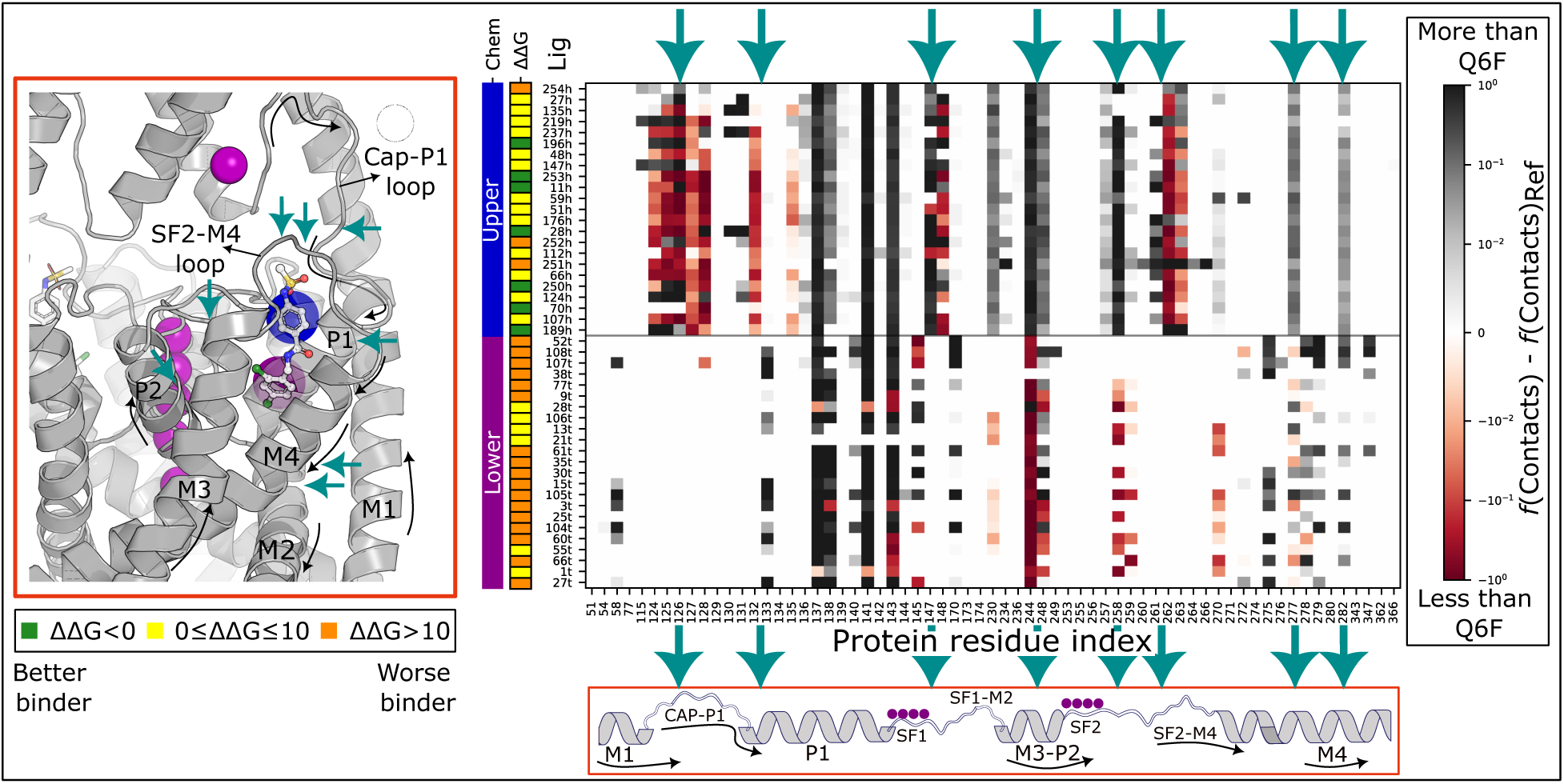
Contact pattern differences of the derivatives in the channel-bound state. The middle panel shows the difference between each Q6F derivative and Q6F itself for an order parameter defined as the fraction of maximal contacts between ligand heavy atoms and pocket heavy atoms within a cutoff of 4.5 Å. The color bar on the right is displayed on a signed logarithmic scale to highlight small changes in contact fractions that may be critical for differences in binding affinity. The bars on the left of the middle panel indicate the chemotype (upper- or lower-ring substitution), as well as a classification into good, medium, or weak binders based on the defined ΔΔ*G* range. The scheme in the lower middle panel maps each protein residue index to a structural region of the channel. Teal arrows highlight key differences in the contact patterns of upper- and lower-ring substitutions.

Upper- and lower-ring substitutions display distinct average contact patterns. Upper-ring substitutions show slightly reduced contacts with the Cap-P1 loop, while contacts with the P1 helix (G137 and T141), N147 in SF1, and residues in M3 and P2 (F230, F244, and T248) are enhanced. Contacts are also strengthened in the SF2-M4 loop around Y257-A259, and in M4 at W277 and L282; while a reduction of contacts is observed at S262-D263 (**Figure 4**).

In contrast, lower-ring substitutions increase contacts with the middle and lower P1 region, particularly F133, G137, and T138. Contacts with N147 remain largely unchanged, whereas interactions with F230, F244, and T248 are predominantly depleted or unaffected. Changes in the SF2-M4 loop are more limited, and M4 displays a mixed contact pattern (**Figures 4 & 5**).

These results indicate that affinity is not determined only by interactions within the amphipathic core of the modulator pocket, but also by contacts with neighboring structural elements: Improved affinity is associated with reduced contacts in the Cap-P1 loop, increased contacts with the SF2-M4 loop (Y257-A259), and reduced contacts farther along the same loop at S262-D263. The F230-F244-T248 region also emerges as an important interaction hotspot, particularly for upper-ring substitutions, together with reinforced contacts to hydrophobic residues in M4 such as W277 and L282. In contrast, increased contacts with the middle P1 helix, including F133, G137, and T138, are associated on average with poorer affinity.

Structurally, the F230-F244-T248 hotspot is part of an aromatic interaction network linking M3 (F230), P2 (F244), the selectivity filter (SF1 F145 and SF2 F254), and P1 (F134) through *π*-*π* stacking. The lower aromatic ring of Q6F fits between F134 and F145, helping preserve this conserved network and, in that way, contributing favorably to affinity (**Figure 5**).

**Figure 5:**
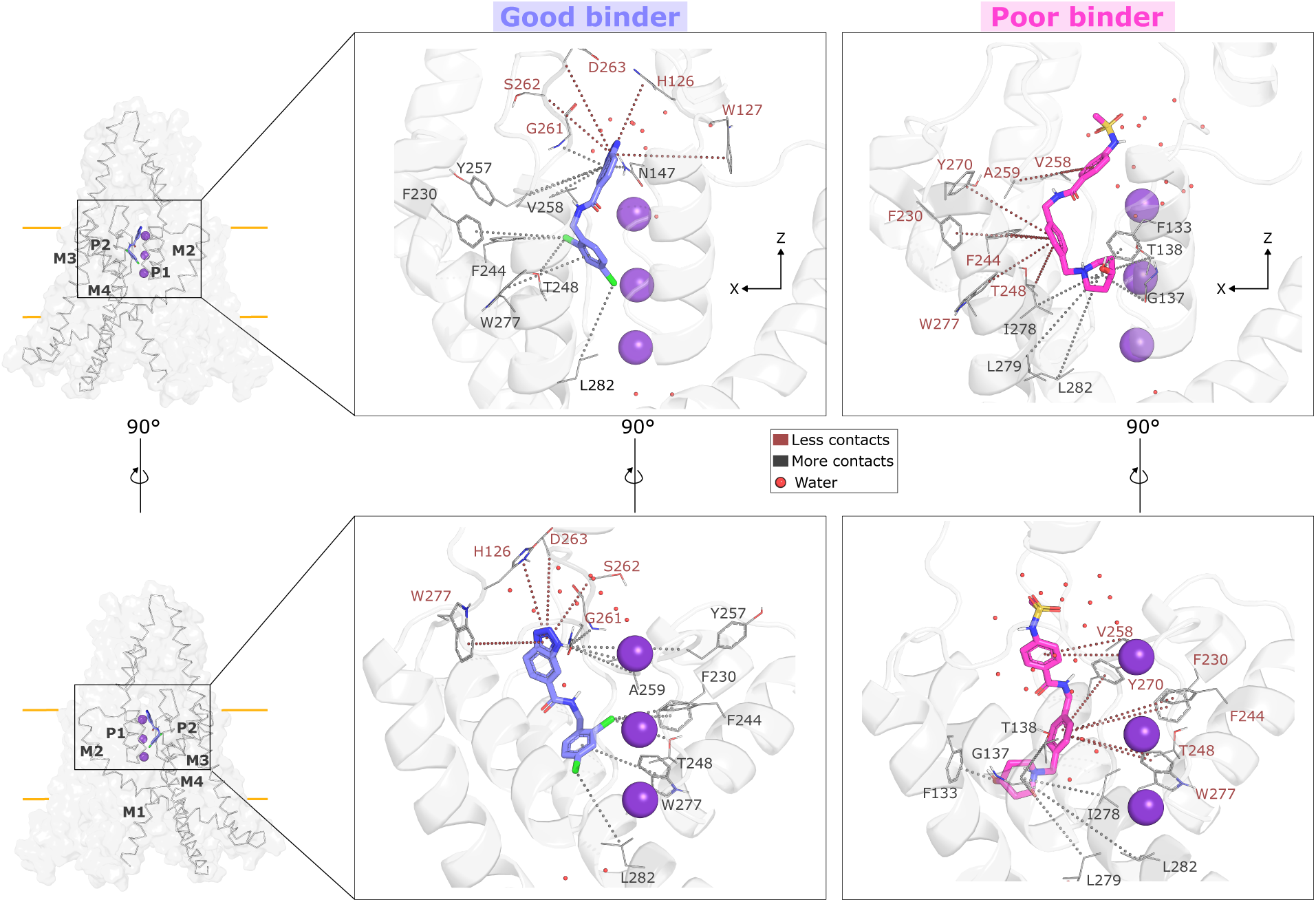
Reshaping of contact patterns in Q6F derivatives. Representative snapshots of a good (253h, ΔΔ*G* = *−*9.36 *±* 0.88 kJ*/*mol) and poor (107t, ΔΔ*G* = 121.12 *±* 10.17 kJ*/*mol) binder relative to the parent compound Q6F. Dotted lines highlight changes in contact patterns within the modulator pocket relative to Q6F, which can be related to the contact pattern matrix shown in Figure 4. Water molecules are shown as small red spheres, whereas K^+^ ions in the SF are shown as purple spheres.

### Contact character, desolvation and conformational restriction upon binding are associated with the asymmetric affinity response

So far, we found that the asymmetric affinity response of the Q6F activator upon chemical perturbations in the TREK1 modulator pocket is associated with differences in contact patterns among the upper and lower pocket. We decided to analyse further properties associated with this asymmetry, that can be more intrinsic to the chemical derivative.

Interestingly, we found that the contact fraction in the ligand-protein interface is higher in contacts between hydrophobic atoms, than between polar ones; this is specially pronounced in the upper-ring substitutions.

Interestingly, we found that the fraction of hydrophobic contacts at the ligand–protein interface is higher than that of polar contacts, particularly for upper-ring substitutions. Further, upon binding, all the chemical derivatives lose, on average, approximately 20-30 water molecules; with the upper-ring substitutions having a greater degree of desolvation (**Figures 6 Panel A-B & S9; Table S2**).

**Figure 6:**
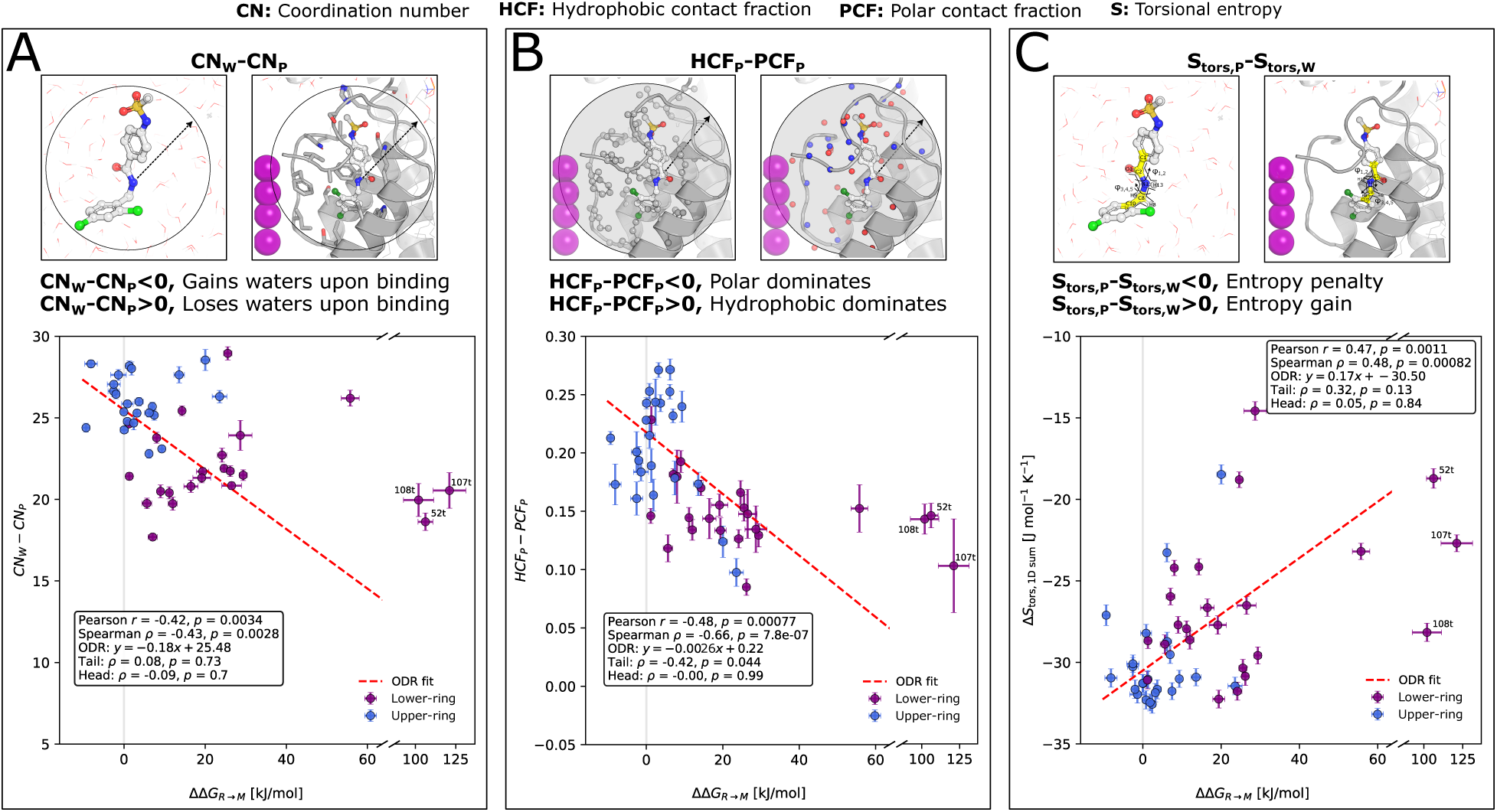
Correlations between Q6F chemical perturbations and desolvation, pocket contacts, and conformational penalization. **Panel A**. Correlation between relative binding free energy and the change in coordination number (CN) upon Q6F perturbation in the solvated (w) versus pocket (p) ensembles. **Panel B**. Correlation between relative binding free energy and the change in hydrophobic versus polar contact fraction in the pocket ensemble. **Panel C**. Correlation between relative binding free energy and the change in entropy associated with the *φ*_1_*_−_*_5_ ligand dihedral distributions in the pocket (p) versus solvated (w) state. The corresponding N-methylacetamide dihedrals are highlighted in yellow in the insets. **All panels**. Correlations were calculated from per-ligand means (N=46) using Pearson’s *r* and Spearman’s *ρ*, with two-sided *p*-values from the *t*-distribution. Spearman’s *ρ* was also calculated separately for the upper- and lower-ring perturbation series (N=23 each). Linear trends were fitted by orthogonal distance regression (ODR), weighting by the inverse squared SEM in both affinity and the corresponding parameter; slope and intercept uncertainties are reported as ODR standard errors.

Finally, we measured a proxy of conformational entropy for the derivatives, that describes the conformation of the N-methylacetamide group linking the upper and lower rings, and we observed that perturbations with better affinity tend to exhibit, on average, a larger entropy penalty upon binding, suggesting stronger conformational restriction in the pocket, whereas weaker binders remain comparatively less constrained. The improvement in binding affinity despite a larger entropic penalty indicates that binding is primarily driven by favorable enthalpic contributions, with conformational restriction in the bound state acting as a secondary thermodynamic factor. This also suggests that less flexible molecules might be even higher in affinity, since do not suffer from conformational restriction upon binding.

It is important to note that this approximation neglects correlations between dihedral angles, as well as contributions from other internal degrees of freedom of the ligand and from the translational, rotational, and desolvation entropy of both the ligand and the protein. Nevertheless, it captures a consistent trend across our chemical series, assigned to the common linker motif shared by all derivatives (**Figures 6 Panel C & S10; Table S3**): An average difference between the low- and high-penalty molecules (*∼*12 J mol*^−^*^1^ K*^−^*^1^), that translates into a -TΔΔS contribution of *∼*3.5 kJ mol*^−^*^1^ to the affinity at 298.15 K, i.e., *∼*1.4 K_b_T (**Table S3**).

### Chemical features associated with the asymmetric affinity response

Finally, we aim to understand the asymmetric response of affinity in the pocket, by simple relationships of physicochemical features of the Q6F derivatives. Thus, we fitted a penalized linear regression model that allows a more straightforward interpretation of how individual descriptors relate to binding affinity.

After subtracting the mean affinity of the corresponding upper- or lower-ring substitution group from each of the N=46 molecules (reducing in that way confounding associated with the Upper/Lower-ring separation), the model recovered a measurable predictive signal: A Leave-one-out cross-validation (LOOCV) Spearman correlation coefficient of *ρ*=0.35, and a coefficient of determination of *Q*^2^=0.57, providing an out-of-sample estimate of the association between descriptor-based predictions and molecular affinities (**Figure S11, Panels A and B**). For comparison, the model trained on the pooled affinities achieved *Q*^2^=0.69, *ρ*=0.54 (**Figure S12**, Panels A–B). Both models’ cross-validated *Q*^2^ significantly exceeded the permutation-derived null distribution (p*≈*0.0033 for each; (**Figure S12**, Panel C)). We note that *Q*^2^ is computed against the total variance of the data, and then is inflated by the few extreme lower-ring compounds. The Spearman *ρ* coefficient is insensitive to this variance magnitude, and is the more conservative description of the model’s ranking performance.

Overall, this analysis indicates that the affinity of lower-ring substitutions correlates with a reduced contribution of features such as molecular surface area from moderately electron-rich atoms, average lipophilicity, the topological distribution of molar refractivity, and favorable patterns of mass dissimilarity across the molecular graph. In contrast, upper-ring substitutions tend to preserve these properties, particularly the contribution of electron-rich aromatic environments to the molecular surface area and a more balanced distribution of lipophilicity.

Likewise, lower-ring substitutions involving saturated nitrogen-containing heterocycles can increase the surface area contributed by slightly positively charged atoms and promote topological clustering of atoms with similar mass or electronegativity. According to the model, both features are associated with weaker affinity. Upper-ring substitutions appear less prone to introducing these detrimental features, particularly because they preserve the aromatic character of the lower ring.

In summary, the consensus model suggests that stronger affinity is associated with increased lipophilicity and specific distributions of weakly polarized, mild electron-rich, and electrotopologically heterogeneous surface, whereas weaker affinity is more strongly associated with saturated nitrogen-containing heterocycles, hydrogen-bond-donor groups, electro-positive surface and increased molecular complexity (**Figures S13 & S14; Table S4**).

## Discussion

Our approach reveals that chemical perturbations within the modulator pocket point toward affinity regions that extend beyond the core of the pocket itself and, interestingly, overlap with regions previously implicated in TREK channel gating. A particularly relevant region is the F230-F244-T248 triad, whose proximity to gating residues reported for TREK1/2 and hERG, together with the putative *π*-*π* stacking network in which such triad is engaged, suggests that the coupling between binding and gating goes beyond a simple hypothesis of amphipathic complementarity between ligand and channel.

We hypothesize that reinforcement of this *π*-*π* network might help improving binding. For example, modifications in the upper-ring of the ligand may promote additional interactions while preserving the lower aromatic ring as a well-packed anchor point within the network. This would be a better strategy than introducing bulky or charged groups in the lower-ring, which tend to hinder affinity. Since endogenous ligands such as lipids are less bulky and more flexible than a small-molecule, they may be able to accommodate themselves within this *π*-*π* stacking, without strongly penalizing the network.

The overlap between the affinity hotspots identified here and previously reported gating residues further supports this interpretation. In TREK1, external acidification has been shown to induce inactivation, likely through a C-type-like mechanism involving H126 in the Cap-P1 loop, Y257 in the SF2-M4 loop, and D263 farther along the same loop.^43^ Under acidic conditions, protonation of H126 has been proposed to promote interaction with D263, inducing conformational changes that propagate toward the filter.^43^ These residues fall within regions that also emerge from our results as important for binding. Thus, an upper-ring substituent may not only improve affinity, but could potentially modulate the relative motion or interaction of these loops, for instance by acting as a physical blocker between them.

A similar overlap is observed around the M3-P2 region. Residues F230, F244, and T248 form an interaction hotspot in our analysis, while previous MD simulations and electrophysiology experiments have suggested that residues in P2, particularly the TREK2 equivalent of T248, transduce activation signals from a distant cytosolic sensor domain to the selectivity filter and regulate inactivation.^44^ In addition, the M3 glutamate network has been shown experimentally to transduce pressure- and temperature-dependent gating cues toward the TREK1 SF.^10^ F230 lies immediately below this network, while residues such as G260 and Y270 are located within the broader SF2-M4 region, where we identify additional affinity hotspots. Together, these observations suggest that the regions acting as anchors for high-affinity modulators may also influence the gating landscape of the channel.

This previous interpretation also allows us to evaluate the initial hypothesis regarding the role of the lower-ring in binding and gating. We initially anticipated that lower-ring substitutions could modulate the S3 carbonyl-flipping mechanism, since this ring lies closest to T141, whose stabilization by Q6F has been proposed to reinforce the S3 backbone conformation.^11^ However, the contact analysis argues against this as a favorable route for improving affinity. Lower-ring substitutions increase contacts with the middle P1 helix, including T138 and G137, but at the cost of interactions involving the F230/F244 *π*-*π* network, and none of these substitutions improve affinity relative to Q6F. Thus, perturbing this ring to influence the carbonyl flip appears to disrupt interactions on which the bound state already depends.

By contrast, upper-ring substitutions preserve the lower aromatic anchor while modifying contacts toward N147 in SF1 and the Cap-P1 and SF2-M4 loops. This allows the Q6F scaffold to be optimized adjacently to the upper SF, making modulation of C-type-like inactivation more possible. At the same time, preserving the lower ring may help stabilize F145 and prevent its carbonyl flip, since such residue forms the S2 binding site of the SF.

The chemical-property analysis supports the same picture. Optimal binders are not extreme in any single descriptor. Instead, aromatic and moderately polar, mildly electron-rich substituents, distributed polarizability across the molecular graph, and amphipathic character are correlated with better affinity; particularly on the upper ring, In contrast, bulky nitrogen-containing saturated heterocycles, strongly localized electropositive patches, overly complex molecular topologies, and substituents that are either too small to satisfy the available contacts or too bulky to fit into the pocket, are generally detrimental. Therefore, our initial hypothesis that bulkier hydrophobic groups in the lower ring and strongly polar or charged groups in the upper ring would systematically improve affinity is only partially supported.

Instead, several contributions appear to follow a balance, or tug-of-war. Moderately polar groups on the upper phenyl ring can increase amphipathic complementarity, but excessively electron-rich surfaces become unfavorable. Likewise, bulky substituents can introduce steric hindrance and localized polarizable heavy-atom patches, whereas groups that are too small fail to satisfy enough interactions within the pocket and can destabilize the binding pose. Improved affinity therefore appears to arise from balancing these competing contributions rather than maximizing any individual chemical feature.

These chemoinformatic observations are consistent with recent experimental results for covalent TREK1 binders.^45^ The Q6F derivative ML336, which carries an acrylamide substitution on the upper ring, retains TREK1 binding and activation, consistent with the idea that aromatic systems displaying polar heteroatoms are well tolerated in this region. Although ML336 forms a covalent bond with S131 under crystallographic conditions, the modified upper ring remains compatible with an active binding mode. A similar behavior is observed for CAT335, a Q6F analogue containing a maleimide substitution that activates the TREK1 S131C mutant and binds covalently to the channel. ^45^ In both cases, substantial modification of the upper ring remains compatible with productive binding.

Another experimentally characterized molecule, C3001a,^16^ also displays activity toward TREK1 and has been proposed from mutagenesis and computational modelling to bind within the same pocket. Although it does not derive directly from the Q6F scaffold, its chemical organization follows a similar pharmacophore: An aromatic, heteroatom-substituted lower moiety and a cyclopentane carboxylate group connected to a phenyl ring through an amide in the upper region. This suggests that hydrophobic contribution from the upper portion of the ligand may compensate for the localized negative charge of the carboxylate and still allow favorable binding.

Finally, we acknowledge that our calculations probe chemical perturbations within the modulator pocket for an activator bound in a single crystallographically derived conductive-state pose. They do not sample the inactivated filter conformation. Therefore, the coupling we infer between pocket occupancy and gating remains an interpretation based on contact geometry rather than a direct measurement of state-dependent affinity. Computing relative affinities against an explicitly inactivated filter model would provide a natural test of the mechanism proposed here, although this remains challenging because no structure has yet been resolved for an activator bound to TREK1 in its inactivated state.

## Conclusions

In this study, we address the question of identifying which regions of the TREK1 modulator pocket carry the affinity signal for better binders, and might be coupled to gating. To this end, we rely on a chemical perturbation of the Q6F activator, and we map such readout onto the pocket structure.

Along these lines, the channel contributions to affinity extend beyond the modulator pocket itself: The hydrophobic lower modulator pocket works as an anchor site, while the upper region (including the Cap-P1 and SF2-M4 loops and the upper SF1 residue N147) plays a more important role in modulating the interaction pattern of activators. This upper region may also serve as a key transducer of gating signals, as suggested by previous experimental results.^10,43^ Moreover, a region encompassing the M3-P2 interface and SF1 configures as an important feature in the middle-lower pocket, in which perturbation by larger groups or point charges induces a penalty arising from both an unfavorable binding pose and a disruption of the *π*-*π* stacking network that we hypothesize is important for controlling SF geometry and, consequently, its gating. A similar logic applies to residue N147, which has previously been identified as a key element in the C-type inactivation of TREK1.^11^

Better Q6F derivatives display a balance of moderate lipophilicity, appropriate molecular size, well-distributed polarizability, and weakly electron-rich surface (as found in polar-substituted aromatic systems) in the upper ring; while avoiding excessively bulky groups (such as nitrogen-containing saturated heterocycles), localized heavy atoms, or electro-positive surfaces that, rather than improving binding enthalpically, give rise to unstable binding modes, particularly at the lower-ring.

Finally, we find that affinity hotspots in the pocket localize around the SF2-M4 loop, the M3-P2 interface, and the upper SF1. The M4 region and the middle pocket act as anchors for hydrophobic phenyl rings, and their perturbation is detrimental for binding. P1 also configures as an anchoring region for the ligand, while certain interactions in the middle of P1 are likewise detrimental for binding, likely by inducing unfavorable binding modes that do not contribute to improved affinity. These hotspots overlap with previous regions identified as important for gating in TREK1. (**Figure 7**).

**Figure 7:**
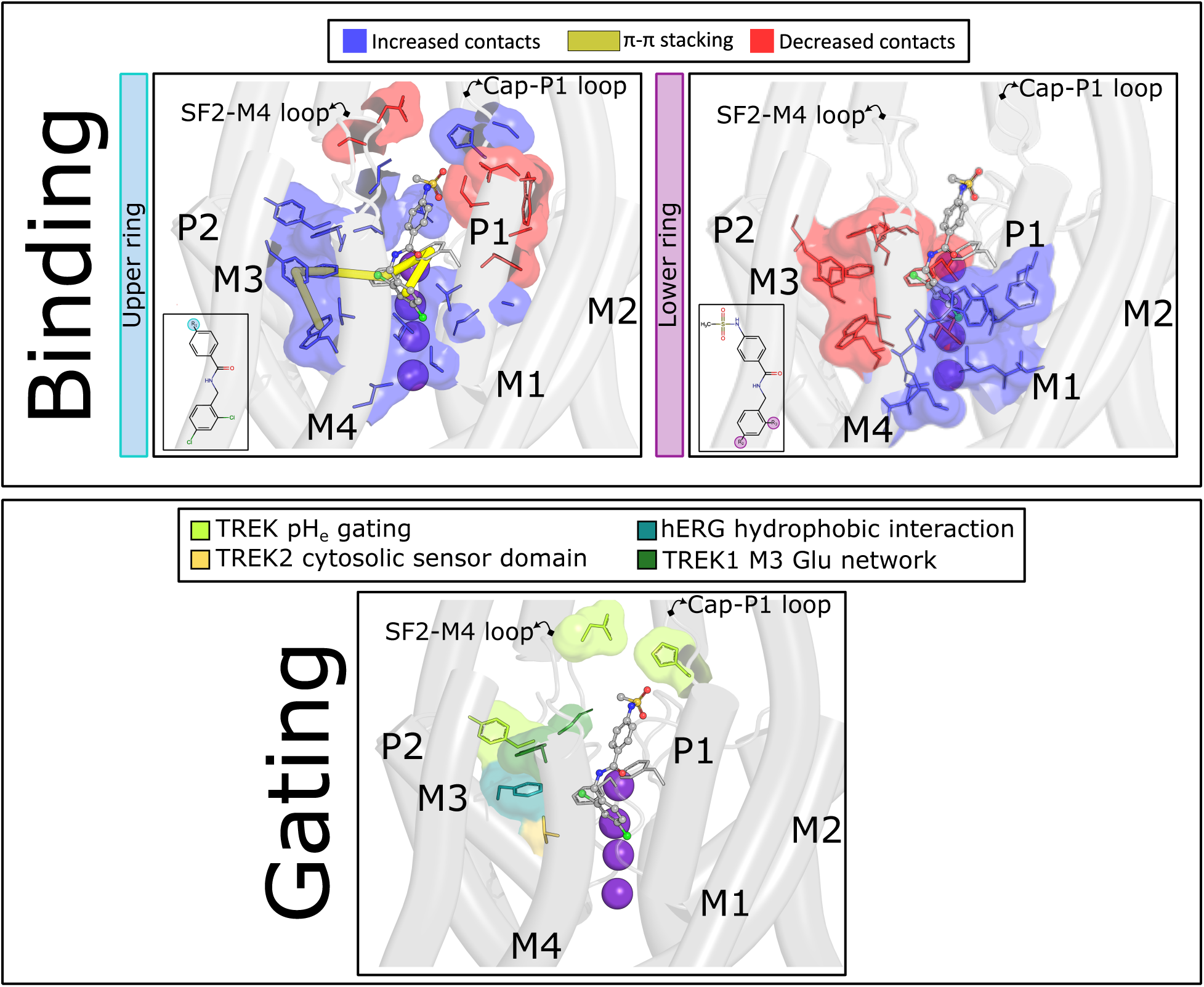
Chemical perturbation map on the TREK1 modulator pocket identifies affinity hotspots overlapping with gating elements. **Binding panel**. Regions where upper- and lower-ring substitutions alter contacts with residues surrounding the modulator pocket. Upper-ring substitutions, generally associated with higher affinity, preserve the *π*-*π* stacking network connecting the M4-M3-P2-P1 helices and SF1 (yellow lines), while forming additional contacts in the upper pocket. Lower-ring substitutions tend to disrupt this network and establish contacts with the lower P1 and M4 helices, potentially introducing unfavorable steric clashes rather than improving affinity. **Gating panel**. Residues previously identified as key elements of SF gating in TREK ^10,43,44^ and hERG ^46^ channels. These residues overlap with regions associated with improved affinity upon upper-ring perturbations.

## Supporting information

Convergence Plots

## Acknowledgement

Edward Mendez-Otalvaro thanks Sudarshan Behera, Chenggong Hui and Carter J. Wilson for discussions about non-equilibrium free energy calculations.

## Supporting Information Available

Raw data and scripts used for this study can be found in: https://github.com/EdwardMendez95/K2P_affinity

