## Supplementary material for "A chemical perturbation scan on a K_2P_ channel modulator pocket identifies affinity hotspots": Convergence Plots

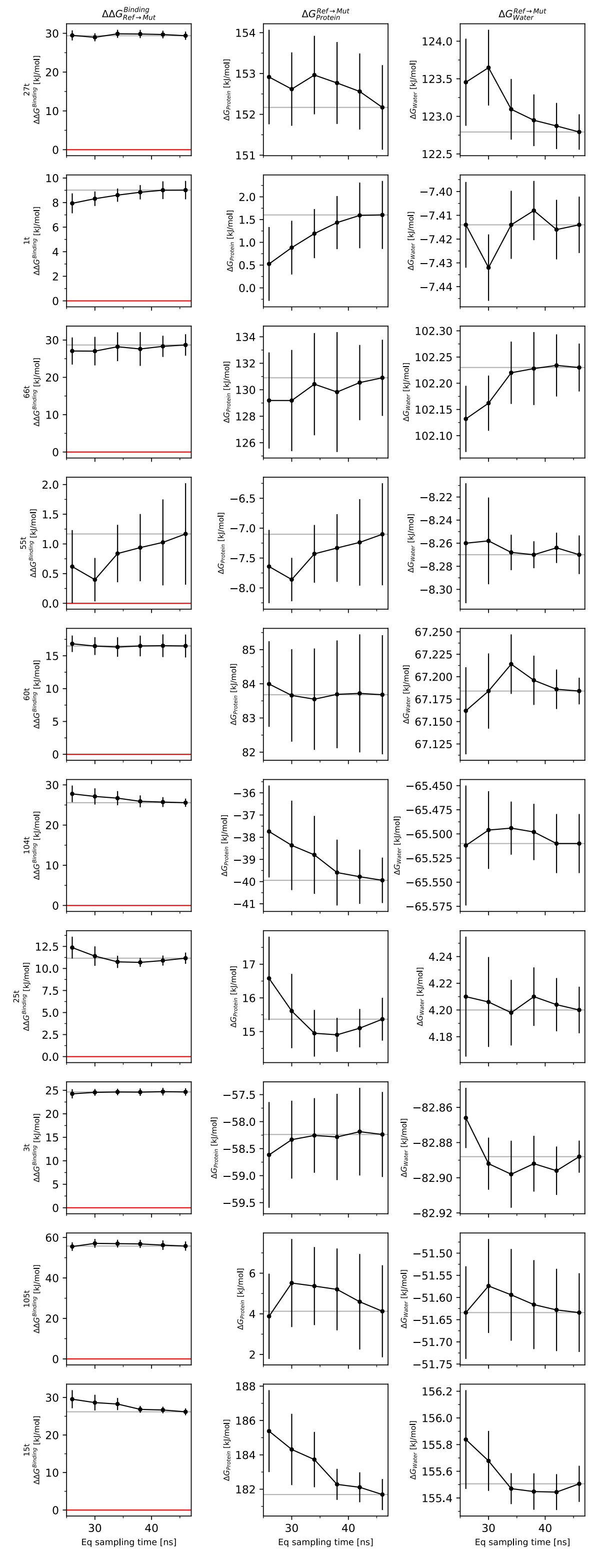


Figure S1A: Convergence of the free-energy estimates as a function of sampling time. Left, middle, and right columns show the relative binding free-energy difference ($\Delta\Delta G$), the protein-leg free-energy estimate ($\Delta G_{\text{protein}}$), and the water-leg free-energy estimate ($\Delta G_{\text{water}}$), respectively. Error bars for $\Delta G_{\text{protein}}$ and $\Delta G_{\text{water}}$ correspond to the analytical SEM across five independent replicates, whereas uncertainties in $\Delta\Delta G$ were obtained by propagation of the corresponding protein- and water-leg errors. Molecule labels are shown to the right of the final column.

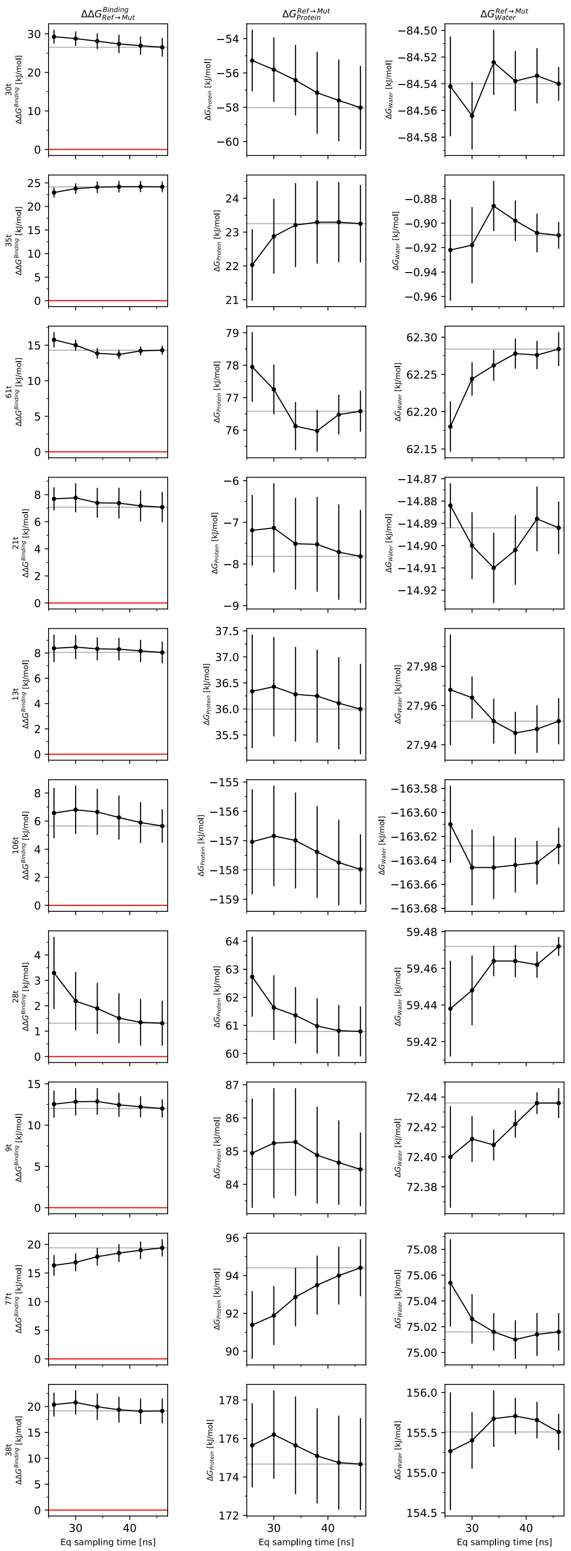


Figure S2A: Convergence of the free-energy estimates as a function of sampling time. Left, middle, and right columns show the relative binding free-energy difference ($\Delta\Delta G$), the protein-leg free-energy estimate ($\Delta G_{\text{protein}}$), and the water-leg free-energy estimate ($\Delta G_{\text{water}}$), respectively. Error bars for $\Delta G_{\text{protein}}$ and $\Delta G_{\text{water}}$ correspond to the analytical SEM across five independent replicates, whereas uncertainties in $\Delta\Delta G$ were obtained by propagation of the corresponding protein- and water-leg errors. Molecule labels are shown to the right of the final column.

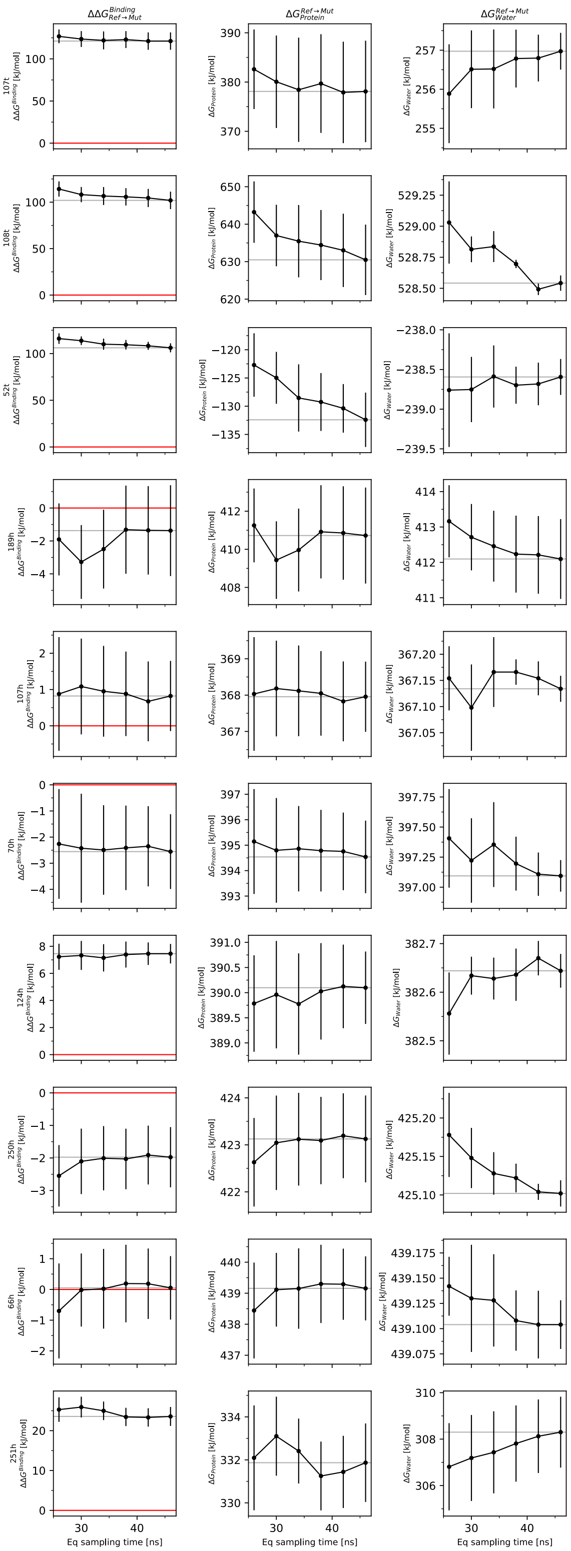


Figure S3A: Convergence of the free-energy estimates as a function of sampling time. Left, middle, and right columns show the relative binding free-energy difference ($\Delta\Delta G$), the protein-leg free-energy estimate ($\Delta G_{\mathrm{protein}}$), and the water-leg free-energy estimate ($\Delta G_{\mathrm{water}}$), respectively. Error bars for $\Delta G_{\mathrm{protein}}$ and $\Delta G_{\mathrm{water}}$ correspond to the analytical SEM across five independent replicates, whereas uncertainties in $\Delta\Delta G$ were obtained by propagation of the corresponding protein- and water-leg errors. Molecule labels are shown to the right of the final column.

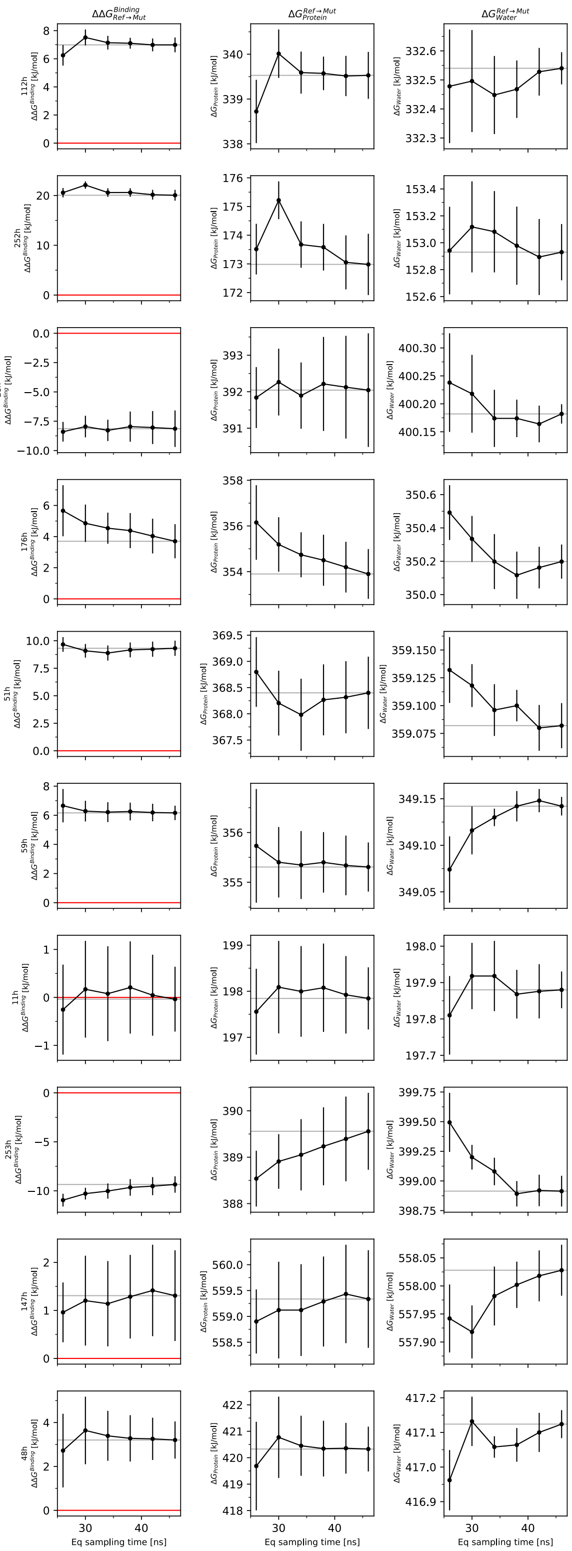


Figure S4A: Convergence of the free-energy estimates as a function of sampling time. Left, middle, and right columns show the relative binding free-energy difference ($\Delta\Delta G$), the protein-leg free-energy estimate ($\Delta G_{\text{protein}}$), and the water-leg free-energy estimate ($\Delta G_{\text{water}}$), respectively. Error bars for $\Delta G_{\text{protein}}$ and $\Delta G_{\text{water}}$ correspond to the analytical SEM across five independent replicates, whereas uncertainties in $\Delta\Delta G$ were obtained by propagation of the corresponding protein- and water-leg errors. Molecule labels are shown to the right of the final column.

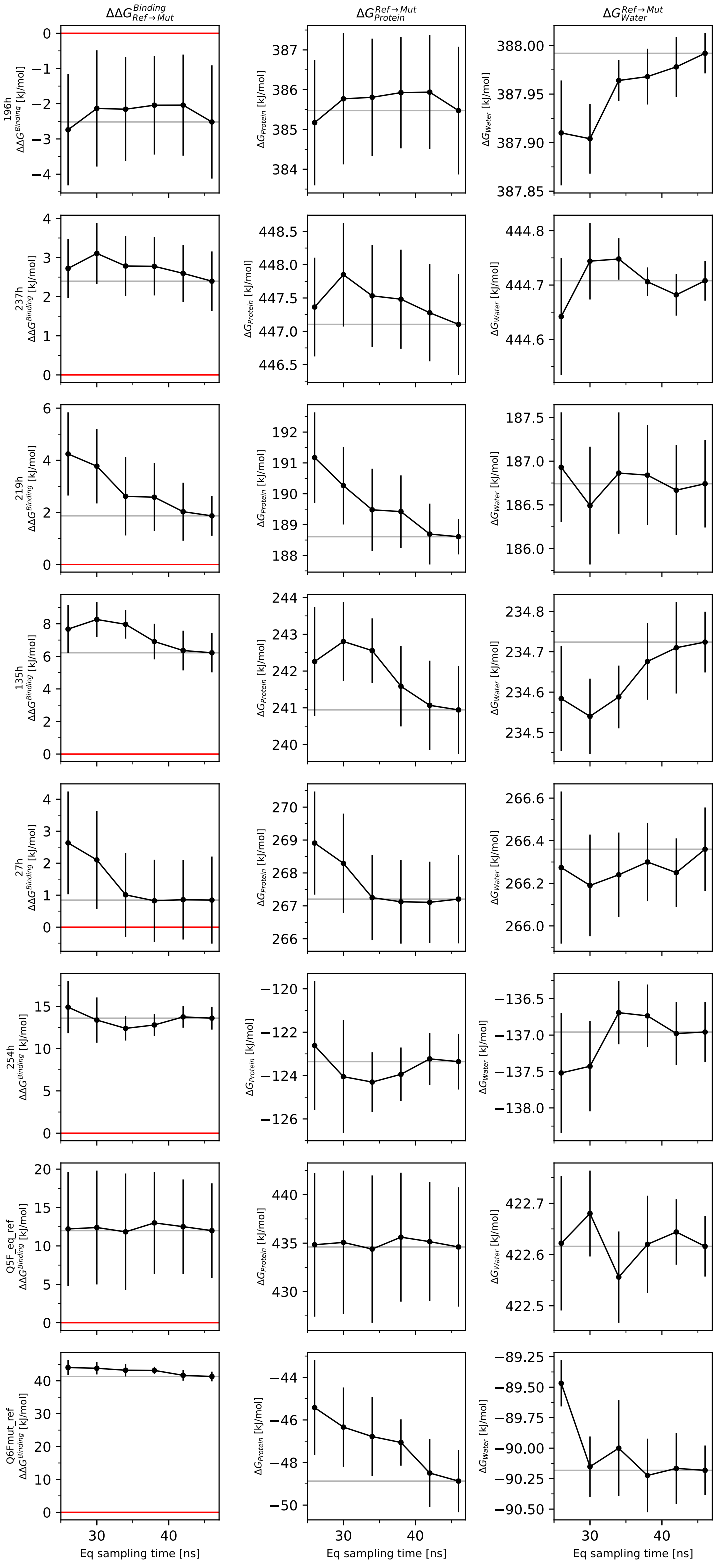


Figure S5A: Convergence of the free-energy estimates as a function of sampling time. Left, middle, and right columns show the relative binding free-energy difference ($\Delta\Delta G$), the protein-leg free-energy estimate ($\Delta G_{\text{protein}}$), and the water-leg free-energy estimate ($\Delta G_{\text{water}}$), respectively. Error bars for $\Delta G_{\text{protein}}$ and $\Delta G_{\text{water}}$ correspond to the analytical SEM across five independent replicates, whereas uncertainties in $\Delta\Delta G$ were obtained by propagation of the corresponding protein- and water-leg errors. Molecule labels are shown to the right of the final column.

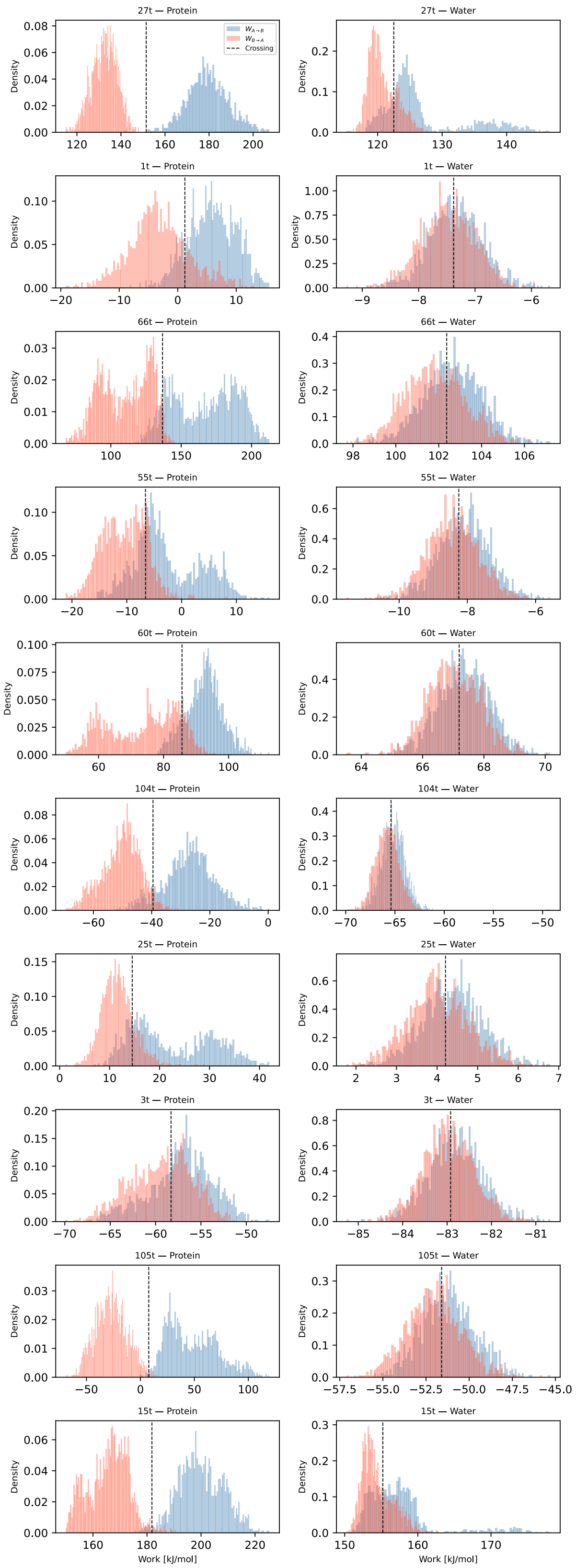


Figure S6A: Overlap of non-equilibrium work distributions for the protein leg (left) and water leg (right). Work values from the five independent replicates were pooled for each transformation. Colors distinguish the forward and reverse work distributions, while the black dotted line marks their crossing point. Molecule labels are shown at the top of each column.

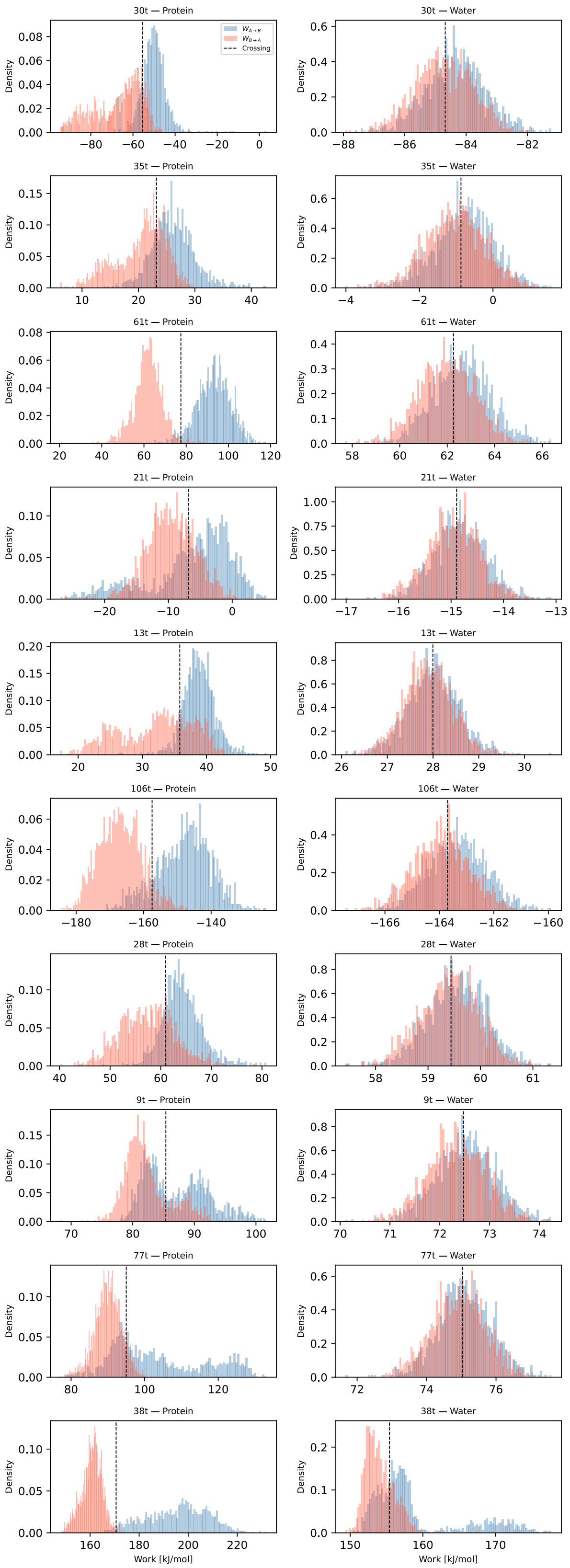


Figure S7A: Overlap of non-equilibrium work distributions for the protein leg (left) and water leg (right). Work values from the five independent replicates were pooled for each transformation. Colors distinguish the forward and reverse work distributions, while the black dotted line marks their crossing point. Molecule labels are shown at the top of each column.

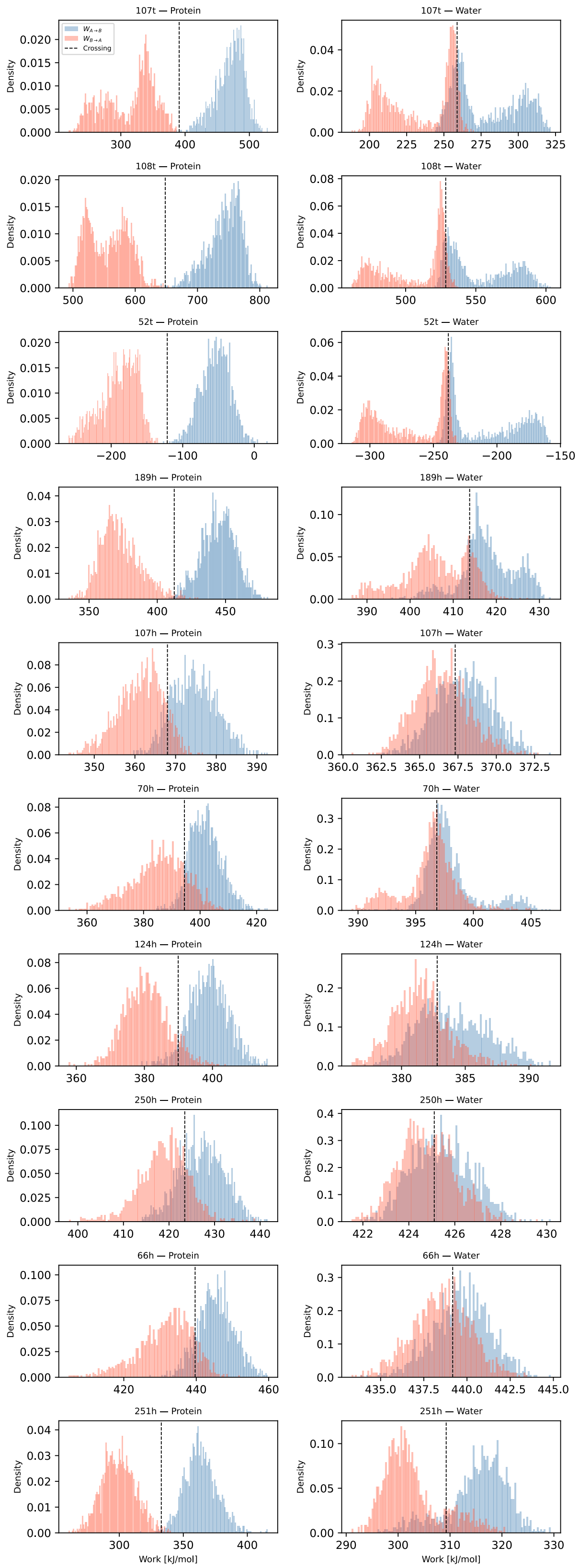


Figure S8A: Overlap of non-equilibrium work distributions for the protein leg (left) and water leg (right). Work values from the five independent replicates were pooled for each transformation. Colors distinguish the forward and reverse work distributions, while the black dotted line marks their crossing point. Molecule labels are shown at the top of each column.

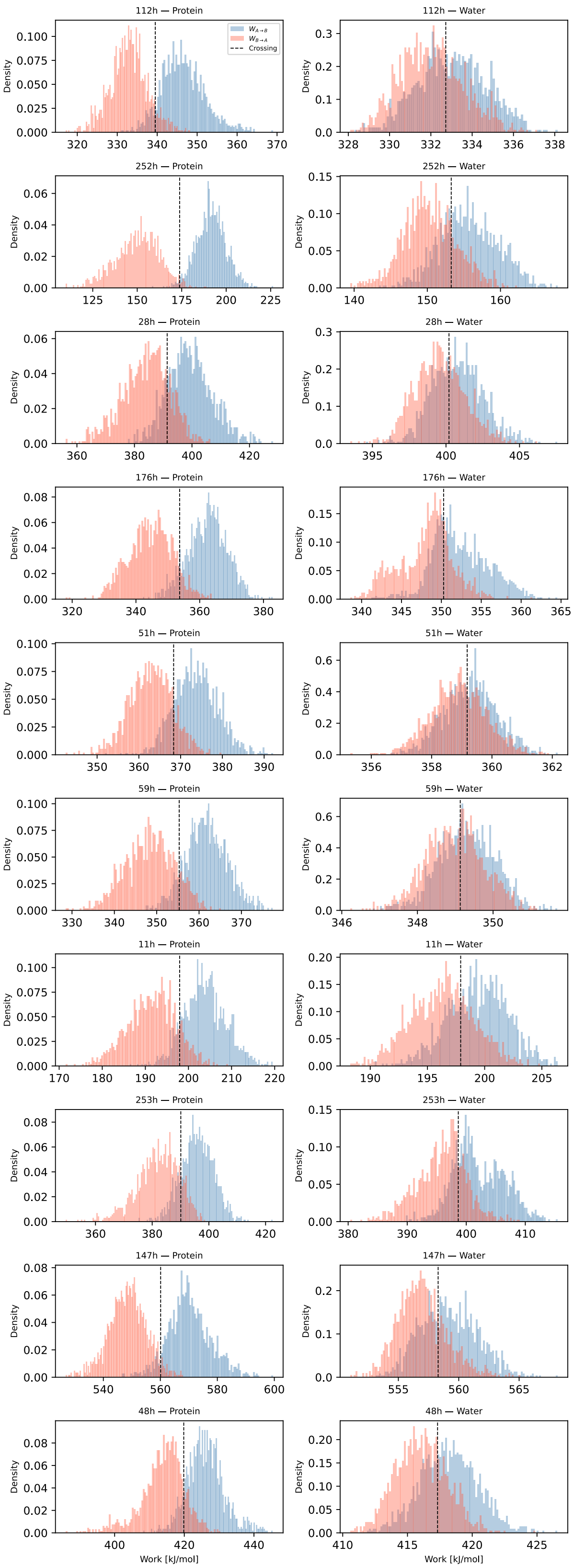


Figure S9A: Overlap of non-equilibrium work distributions for the protein leg (left) and water leg (right). Work values from the five independent replicates were pooled for each transformation. Colors distinguish the forward and reverse work distributions, while the black dotted line marks their crossing point. Molecule labels are shown at the top of each column.

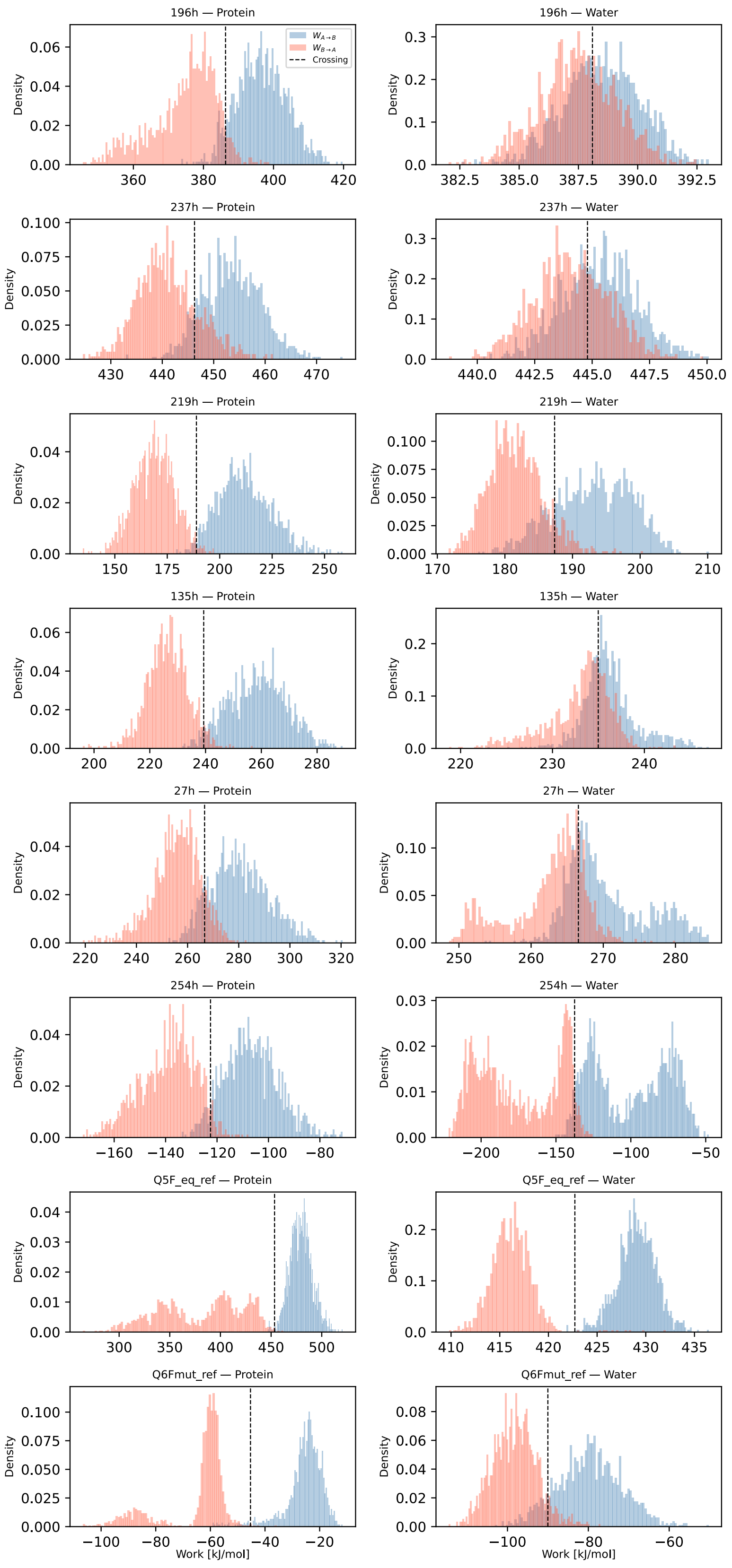


Figure S10A: Overlap of non-equilibrium work distributions for the protein leg (left) and water leg (right). Work values from the five independent replicates were pooled for each transformation. Colors distinguish the forward and reverse work distributions, while the black dotted line marks their crossing point. Molecule labels are shown at the top of each column.
