## Supplementary Information for "A chemical perturbation scan on a K_2P_ channel modulator pocket identifies affinity hotspots"

### Supplementary material

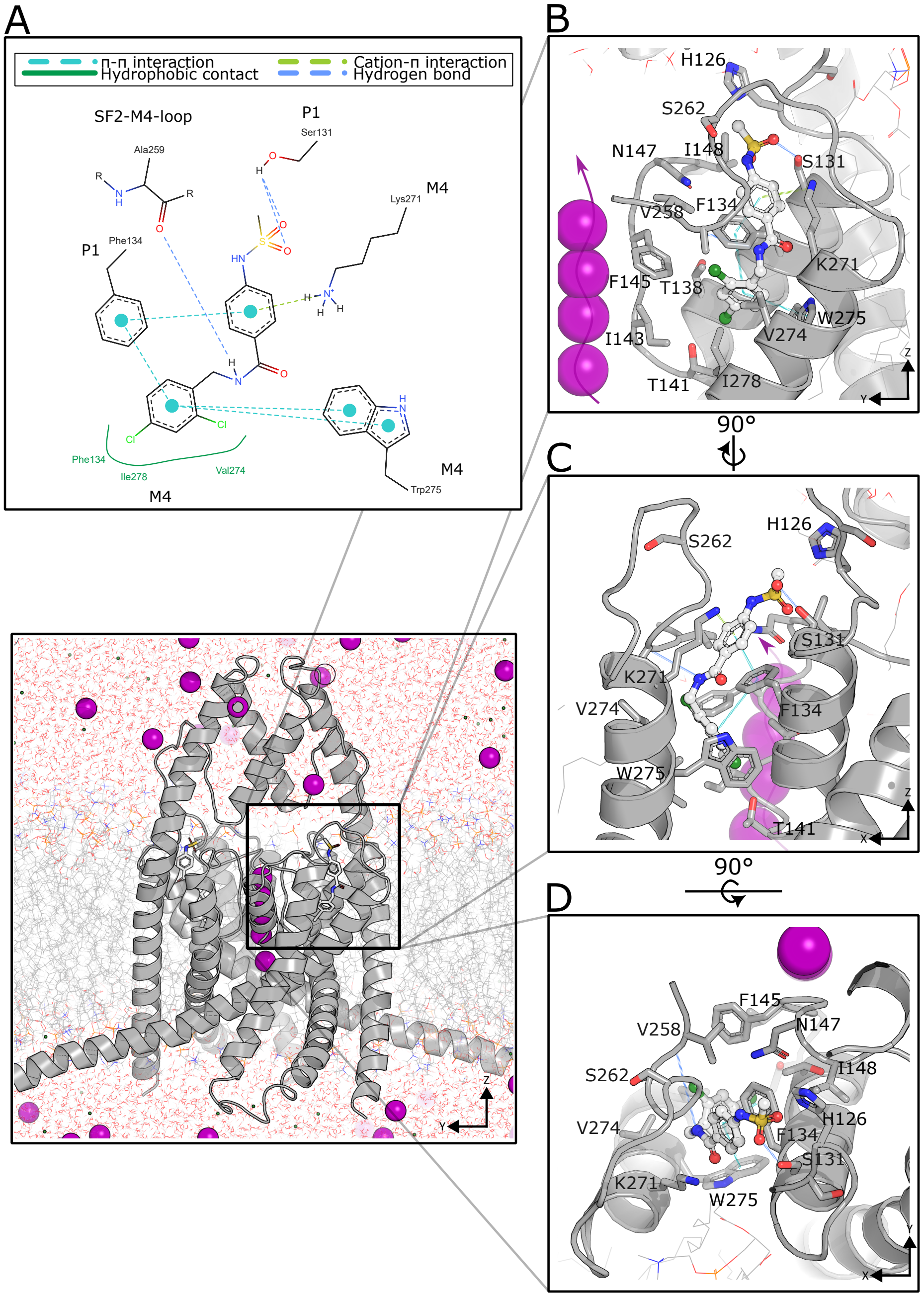


Figure S1: **TREK1 and its modulator pocket. Panel A**: Two-dimensional representation[2] of the main interactions of Q6F (chain A) within the binding site (modulator pocket), based on the x-ray TREK1 holo structure (Protein Data Bank PDB code: 6CQ8). Different types of non-covalent interactions are highlighted. **Panel B**: Zoomed view in the *zy* plane of the Q6F binding pose within the pocket, based on the PDB model (embedded in a solvated membrane simulation box). Key residues belonging to the P1 helix, M4 helix, and SF1 region are labeled. **Panels C-D**: Same view as in Panel B, but shown in the *zx* and *xy* planes, respectively.

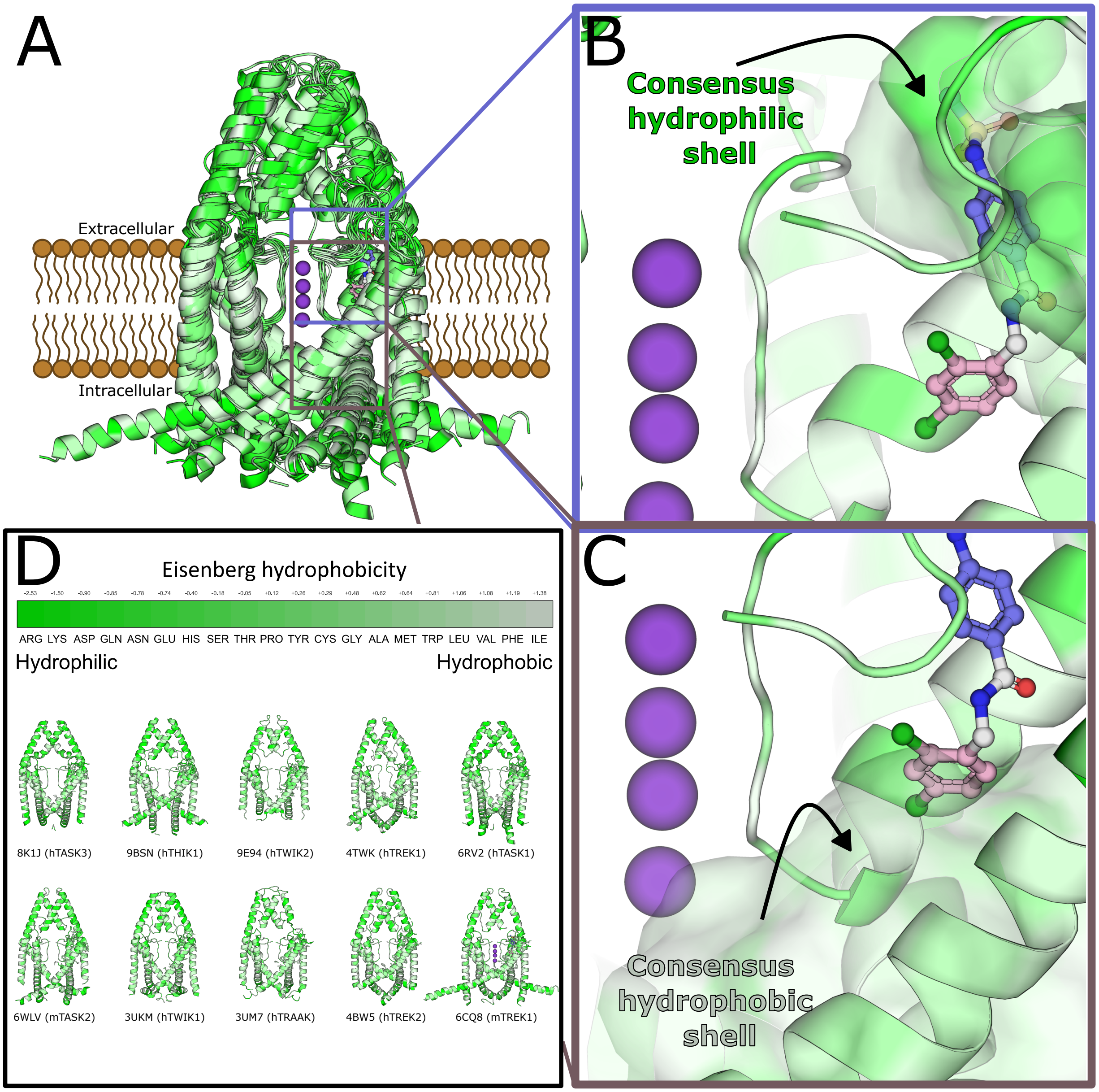


Figure S2: **Hypothesized ligand and channel features contributing to affinity in the TREK1 modulator pocket. Panel A**. Structural superposition of experimentally resolved $K_{2P}$ channel structures aligned around the transmembrane region and SF. Q6F is shown as sticks in the modulator pocket, while $K^+$ ions occupying the SF are represented as purple spheres. Blue and pink boxes indicate the upper and lower regions of the modulator pocket analyzed in Panels B and C, respectively. **Panel B**. Enlarged view of the upper region of the modulator pocket. The upper aromatic ring of Q6F is shown in blue, while the transparent surface represents the consensus hydrophilic environment formed by residues from the aligned $K_{2P}$ structures. **Panel C**. Enlarged view of the lower region of the modulator pocket. The lower aromatic ring of Q6F is shown in pink, while the transparent surface represents the consensus hydrophobic environment formed by residues from the aligned $K_{2P}$ structures. **Panel D**. Eisenberg hydrophobicity scale[1] used to color the cartoon and side-chain representations of the aligned $K_{2P}$ structures. The experimentally resolved channel structures included in the comparison are shown below the scale, with side chains lining the corresponding modulator-pocket regions represented as sticks. The prefixes h and m indicate human and mouse channels, respectively. Together, the structural comparison reveals a conserved amphipathic organization of the modulator pocket, with a predominantly hydrophilic upper region and a more hydrophobic lower region. We hypothesize that this profile contributes to ligand binding by favoring complementary interactions with chemically distinct moieties of bound modulators.

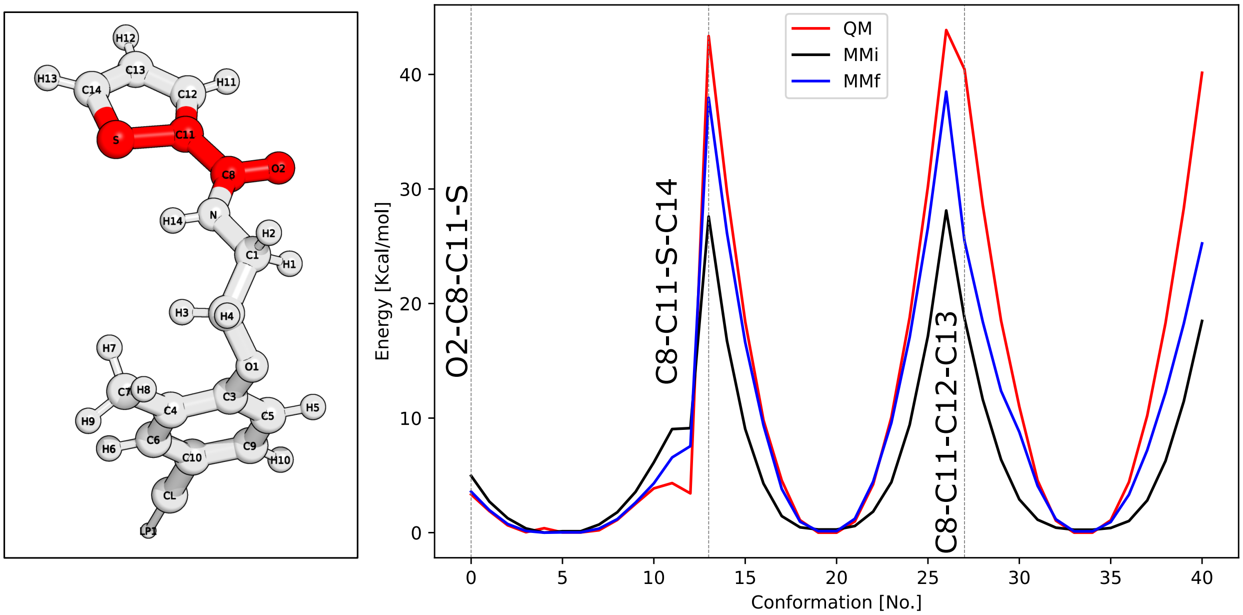


Figure S3: Torsional parameter optimization of high-penalty dihedrals around the thiophene ring of Q5F. The QM energy profile was used as a reference to optimize the initial molecular-mechanics profile (MMi) toward the final molecular-mechanics profile (MMf) by refining the corresponding torsional parameters. See Methodology below.

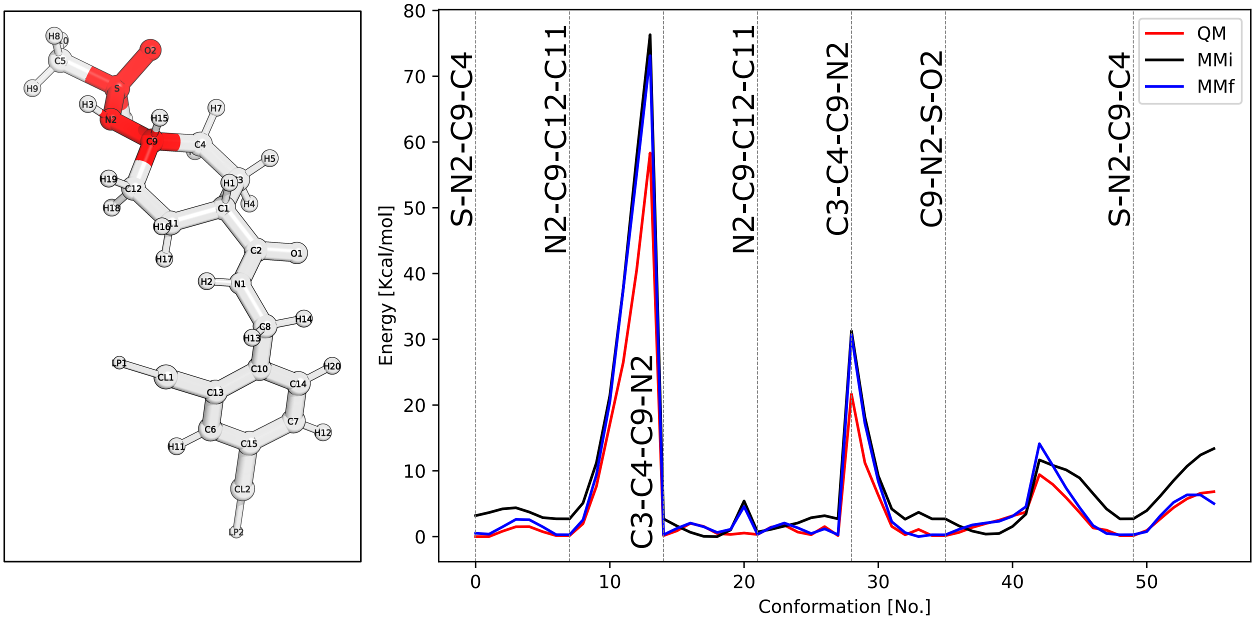


Figure S4: Torsional parameter optimization of high-penalty dihedrals around the cyclohexylsulfonamide moiety of Q6Fa. The QM energy profile was used as a reference to optimize the initial molecular-mechanics profile (MMi) toward the final molecular-mechanics profile (MMf) by refining the corresponding torsional parameters. See Methodology below.

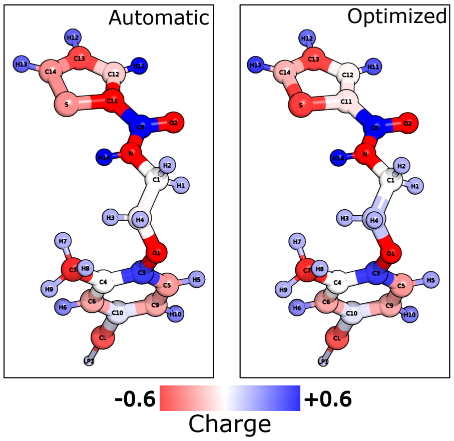


Figure S5: Partial-charge optimization around the thiophene ring of Q5F.

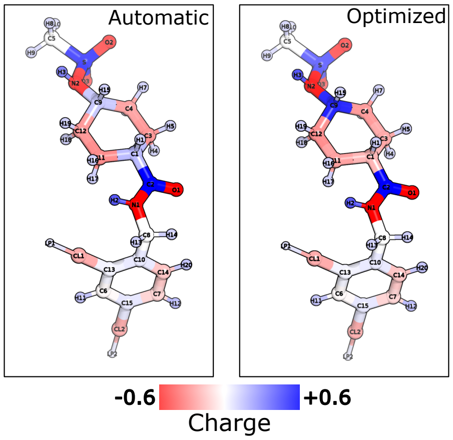


Figure S6: Partial-charge optimization around the cyclohexylsulfonamide moiety of Q6Fa.

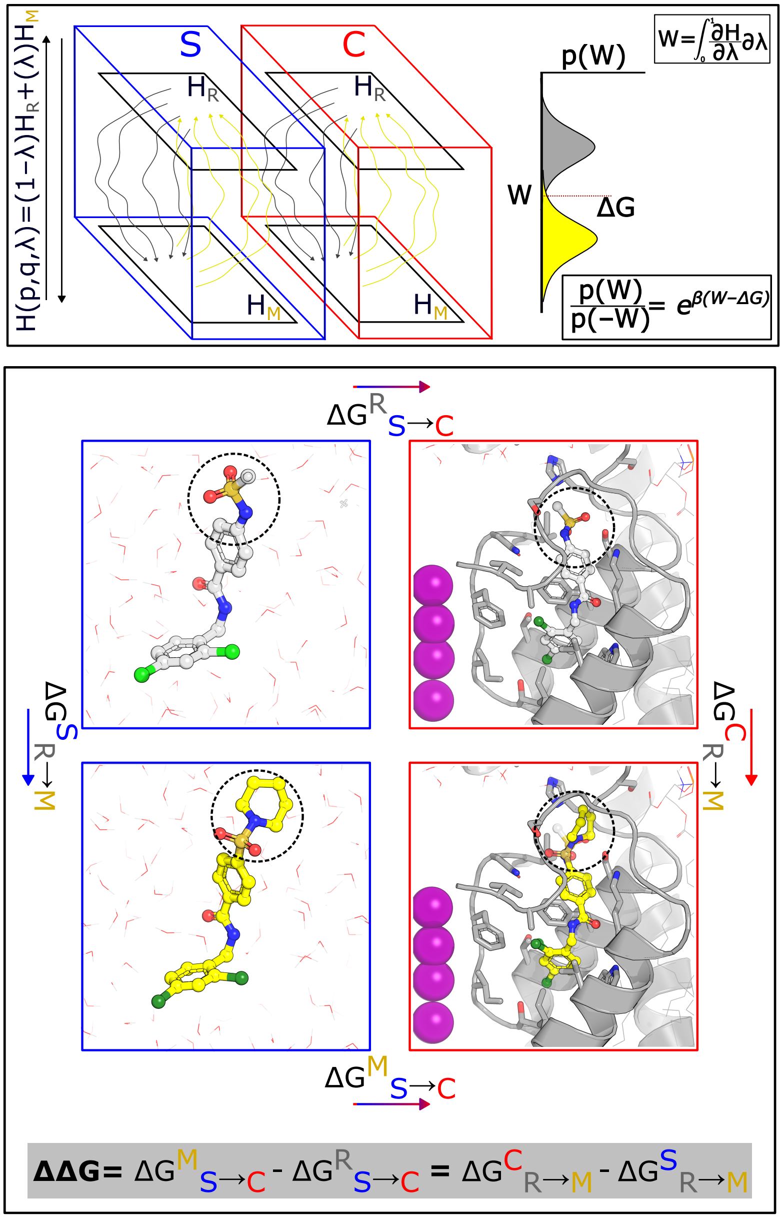


Figure S7: **Estimation of relative binding free energy differences between Q6F and its derivatives. Upper panel**: The relative binding free energy difference is estimated as the difference between two free energy changes computed in the channel (C) and solvent (S) phases. In each phase, the free energy difference between the Hamiltonians of the reference ligand (R, Q6F) and the modified ligand (M, Q6F derivative) is estimated. This is achieved by introducing an alchemical (un-physical) transformation in which the Hamiltonian is interpolated using a coupling parameter $\lambda$ between the two end states. By performing multiple forward and reverse non-equilibrium transitions and recording the work associated with the $\lambda$ derivative, one obtains work distributions whose overlap allows estimation of the free energy difference, as described by the Crooks fluctuation theorem.[47] **Lower panel**: Thermodynamic cycle used to estimate the relative binding free energy difference between Q6F and its derivatives. The quantity of interest corresponds to the difference in binding free energy between the modified ligand (M) and the reference ligand (R) (horizontal branches), which is computationally expensive to evaluate. Instead, the vertical branches (corresponding to the alchemical transformation from R to M in both solvent (S) and channel (C) environments) are computed, as they are more computationally feasible. Since free energy is a state function, the total free energy change around the cycle is zero, and both routes converge to the same result.

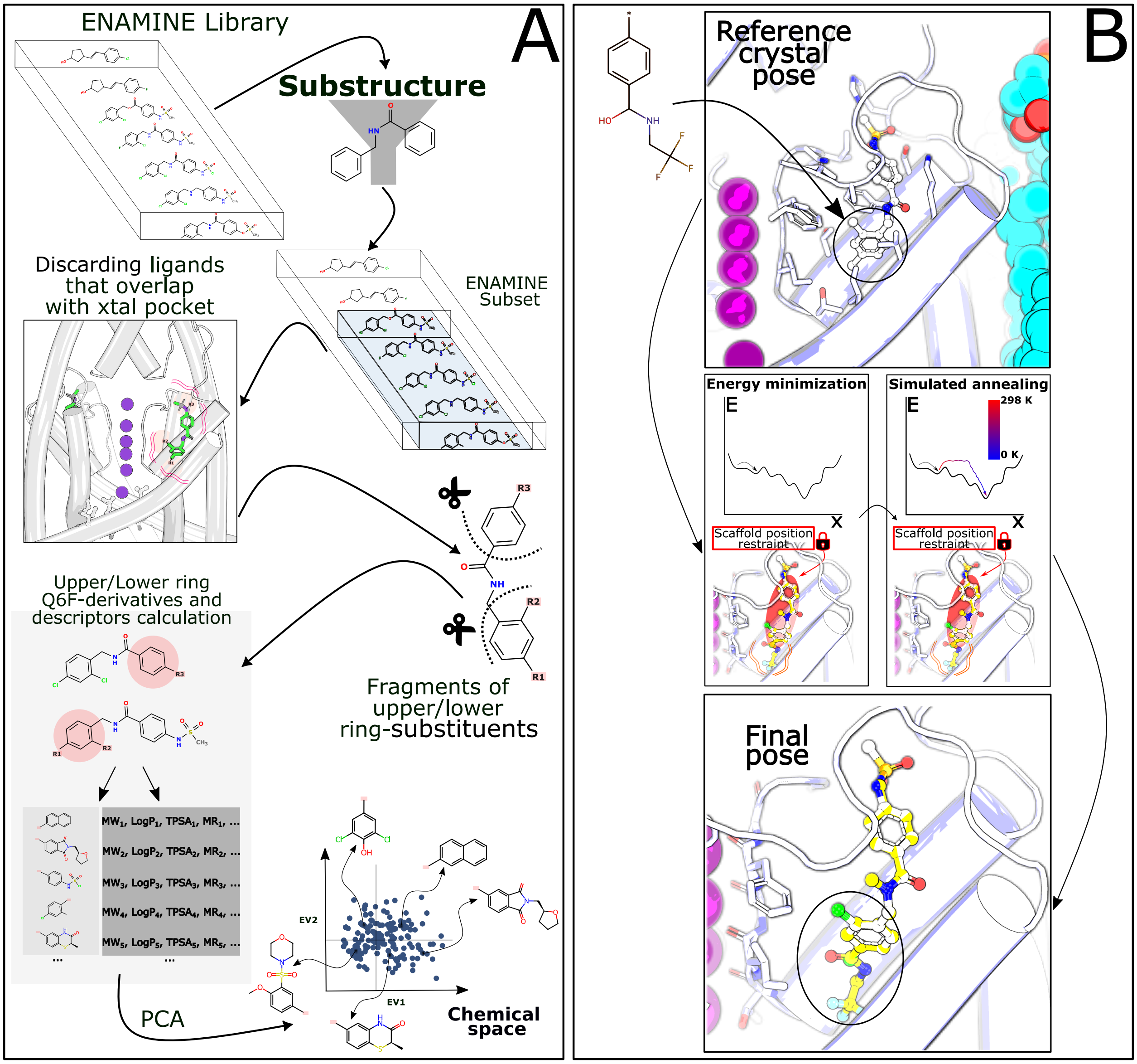


Figure S8: **Protocol for chemical library construction and derivative pose generation. Panel A**: Starting from a database of purchasable compounds (ENAMINE), molecules displaying the N-benzoylbenzylamine scaffold and not sterically overlapping with the binding pocket are selected. Their chemical substituents are then isolated by fragmenting the molecules into upper and lower phenyl-ring fragments. These substituents are subsequently mapped onto the Q6F scaffold, either at the upper or lower ring, and molecular descriptors for the resulting derivatives are computed. The matrix of derivatives versus molecular descriptors is then projected into a low-dimensional space using a Principal Component Analysis (PCA), which captures the main variance of the library. **Panel B**: Once a derivative is selected, its three-dimensional coordinates are placed into the modulator pocket using the N-benzoylbenzylamine scaffold as anchor points. The newly introduced substituents are then energy-minimized while the rest of the system is restrained, followed by a gradual freezing procedure (mimicking cryogenic conditions), producing a final binding pose used for subsequent calculations.

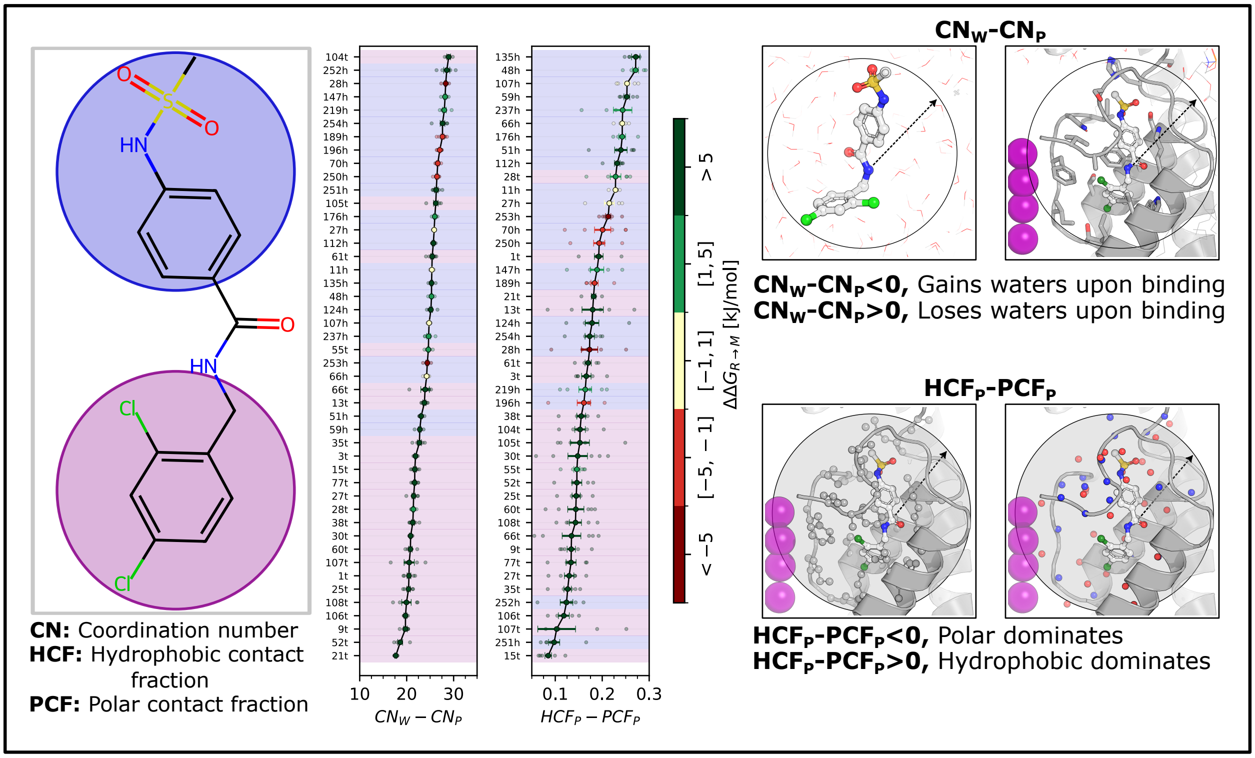


Figure S9: **Correlations between chemical perturbations and interactions with the protein and solvent.** Changes in the coordination number (CN) upon Q6F chemical modification in the solvent (water, w) ensemble; and in the hydrophobic contact fraction (HCF) versus polar contact fraction (PCF) upon Q6F chemical modification in the channel (protein, p) ensemble. The CN is defined as the number of water oxygen atoms within 3.5 Å of any ligand heavy atom. The HCF is defined as the number of protein carbon atoms within 4.5 Å of any ligand heavy atom, while the PCF corresponds to protein nitrogen or oxygen atoms within 4.5 Å of any ligand heavy atom. Markers in the plot are colored according to the relative binding free energy difference, as indicated by the color bar, while the background color (pastel blue or light pink) denotes whether the derivative corresponds to an upper- or lower-ring substitution.

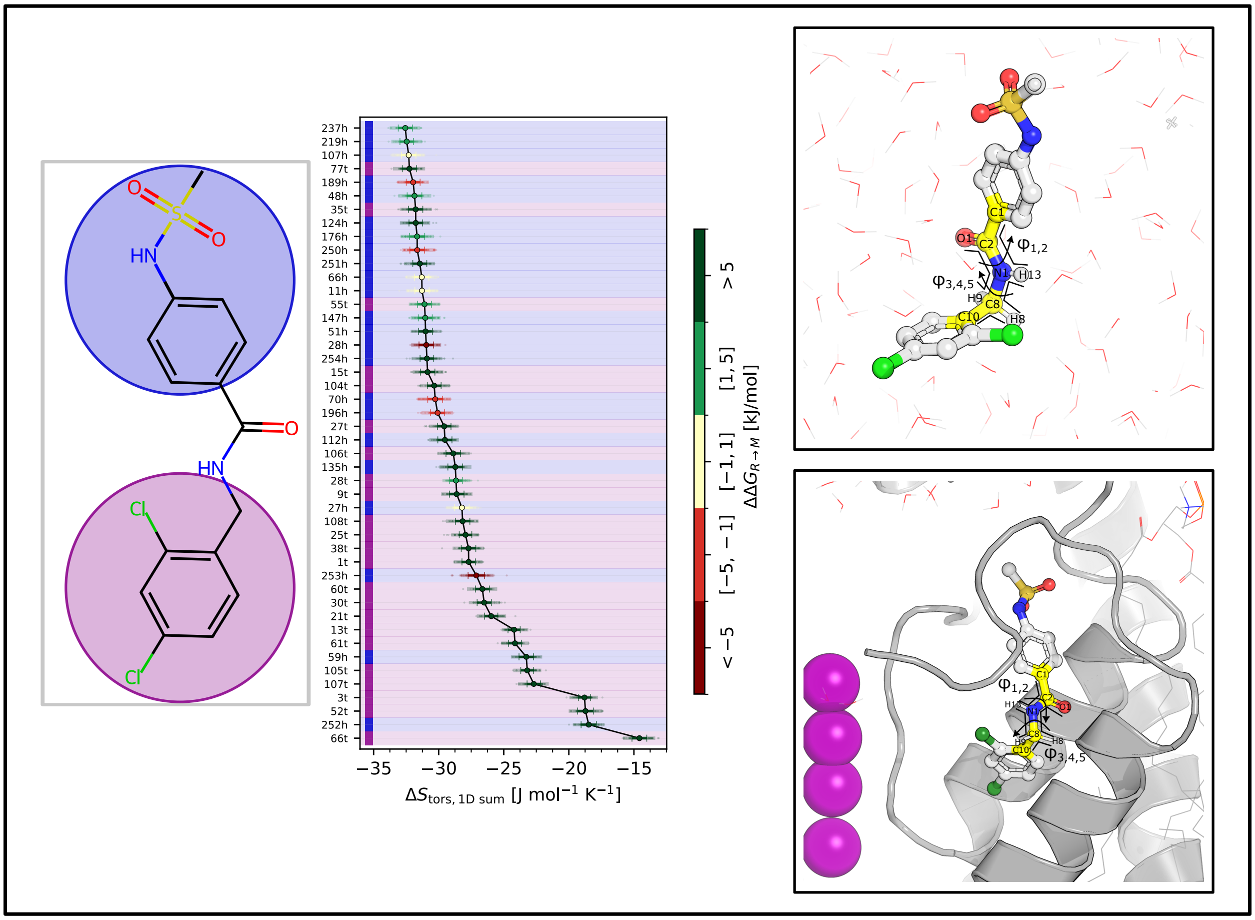

Figure S10: **Conformational entropy of the upper–lower phenyl-ring linker correlates with binding affinity differences.** Difference in entropy associated with the $\varphi_{1-5}$ dihedral angle distributions of the ligand in the channel versus the solvated state. The corresponding dihedral angles are highlighted in yellow in the right insets, showing the molecule in both the bound (channel) and solvated states. Markers in the middle panel are colored according to the relative change in binding affinity, as indicated by the color bar, while the background color denotes whether the substitution occurs on the upper or lower ring. More negative values indicate a larger entropy penalty upon binding.

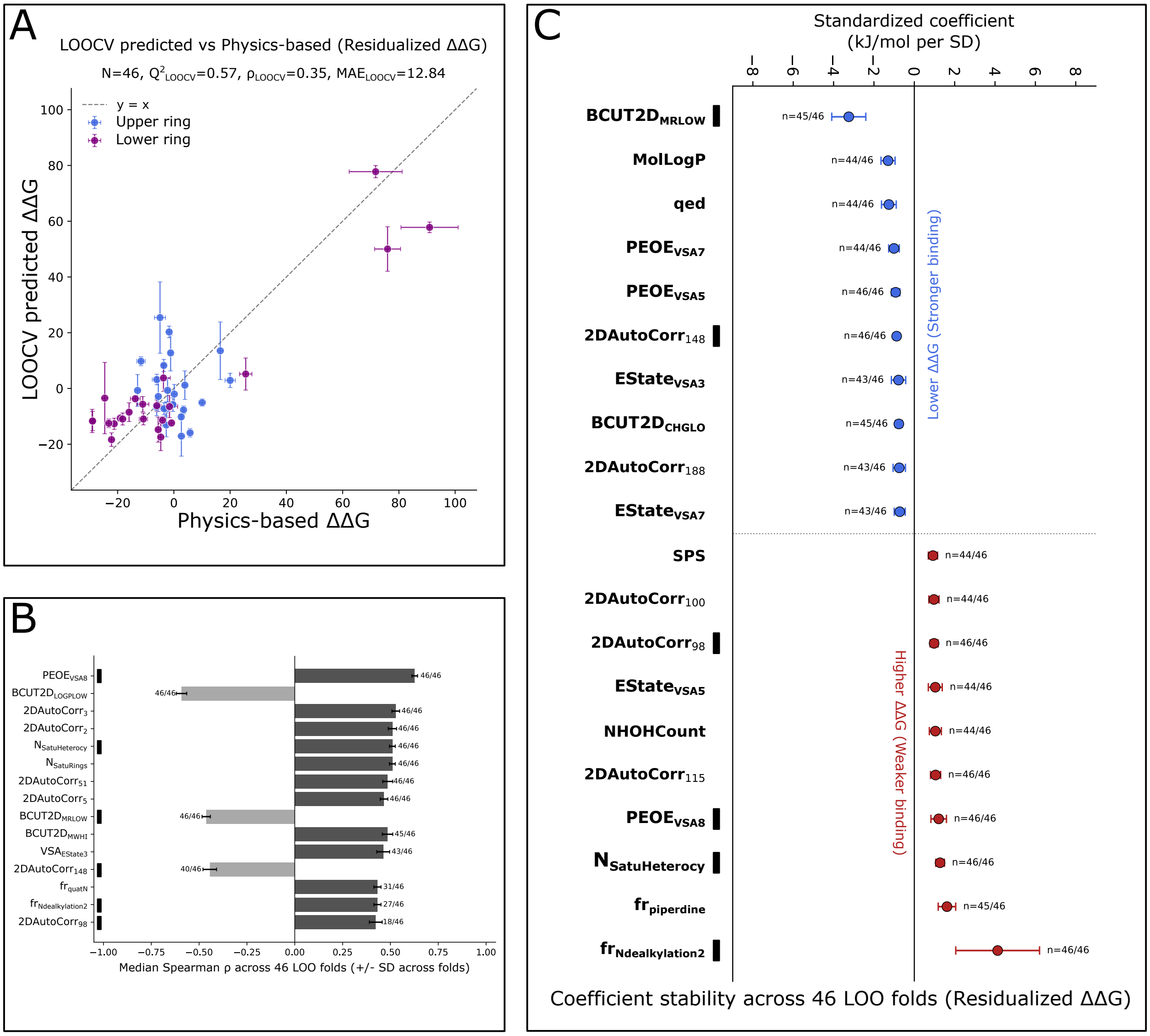


Figure S11: **Chemical interpretation of the QSAR model for the Q6F derivatives. Panel A.** Predicted versus Physics-based affinities for the 46 Q6F derivatives using the residualized ElasticNet model. Model performance was evaluated using nested leave-one-out cross-validation (LOOCV), with one molecule held out in each outer fold and ElasticNet hyperparameters optimized by five-fold cross-validation within the corresponding training set. Vertical error bars represent the standard deviation of the predicted value across repeated nested-LOOCV calculations using different inner-cross-validation partitions. Horizontal error bars correspond to the standard error associated with the physics-based $\Delta\Delta G$ estimates. **Panel B**. Stability of the univariate association between molecular descriptors and residualized affinity. In each of the 46 leave-one-out folds, Spearman correlations were recalculated using the remaining 45 molecules and descriptors were ranked according to $|\rho|$. The 15 descriptors shown correspond to those most consistently ranked among the top 15 across the 46 folds. Bars indicate the median Spearman correlation coefficient across folds, error bars represent the corresponding standard deviation, and the fractions indicate the number of folds in which each descriptor was included among the top 15. **Panel C**. Stability of molecular-descriptor coefficients in the residualized ElasticNet model across the 46 outer LOOCV folds. Points represent the mean standardized ElasticNet coefficient across folds and error bars indicate the corresponding standard deviation, with folds in which a descriptor was not selected contributing a coefficient of zero. Fractions next to each descriptor indicate the number of LOOCV folds in which it received a non-zero coefficient. Black squares identify descriptors additionally selected for chemical interpretation based on their univariate Spearman association with affinity ($|\rho| > 0.5$, $p < 0.05$); the same markers are shown for the corresponding descriptors in Panel B. See the main text for their chemical interpretation.

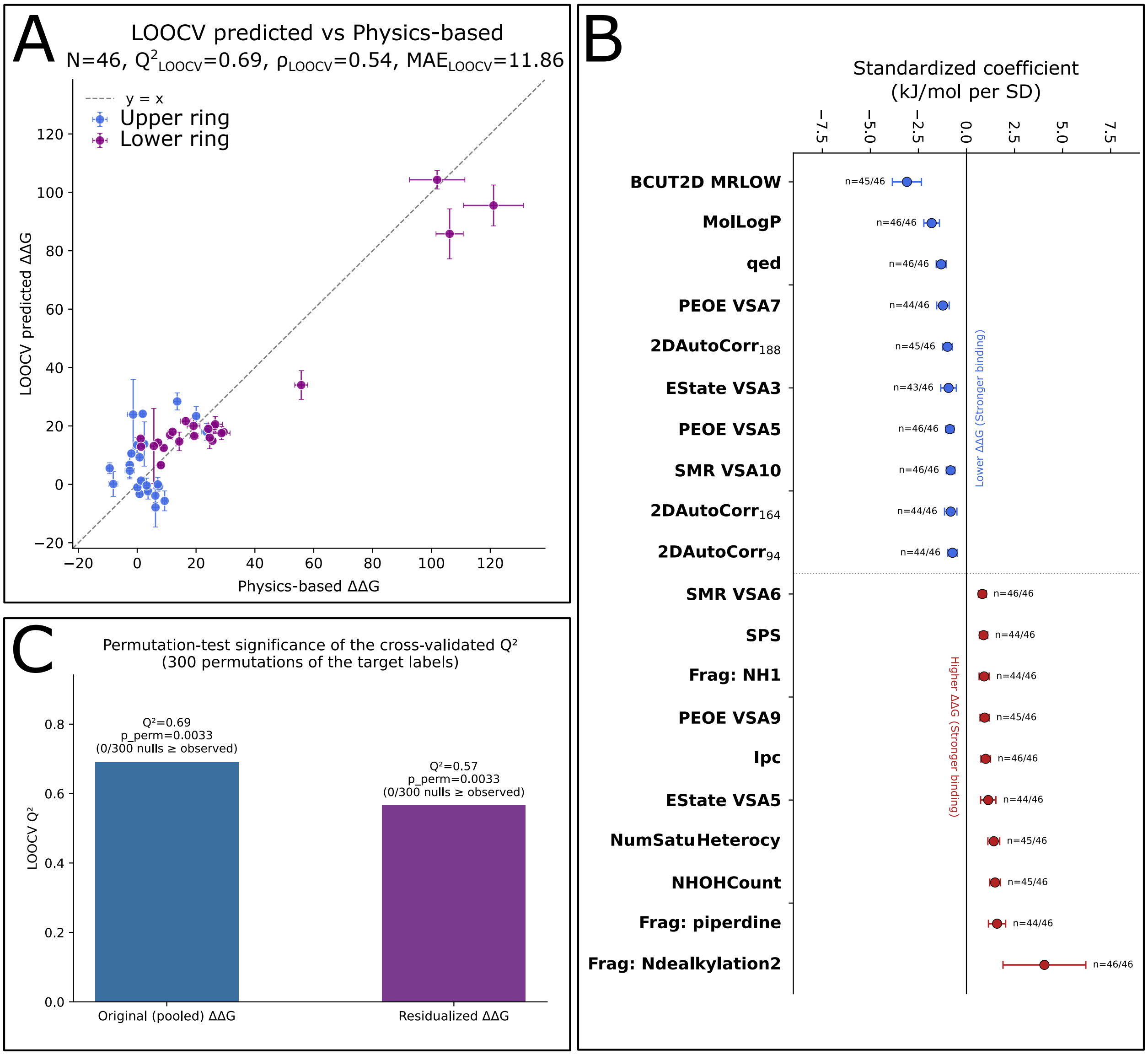


Figure S12: **Performance and significance of the ElasticNet model fitted to the original (pooled) $\Delta\Delta G$ target**. **Panel A** Leave-one-out cross-validated (LOOCV) predicted vs. physics-based $\Delta\Delta G$ for the pooled, non-residualized affinity model. Points are colored by chemotype (Upper-ring, blue; Lower-ring, purple); vertical error bars give the SD of each molecule's predicted value across repeated nested-LOOCV runs using different inner-CV partitions. **Panel B** Standardized ElasticNet coefficients for descriptors with nonzero weight in $\geq 1$ of the 46 leave-one-out folds, grouped by direction of effect (blue: associated with lower $\Delta\Delta G$ (stronger binding); red: associated with higher $\Delta\Delta G$ (weaker binding). Points are the mean coefficient across folds $\pm$ SD; the fraction of folds in which each descriptor was selected is shown over markers. **Panel C**. Permutation-test significance of the cross-validated $Q^2$ for the pooled and region-residualized models. Bars show the observed LOOCV $Q^2$ for each target; annotations give the number of the 300 label-permuted null models (out of 300) that matched or exceeded the observed $Q^2$, and the resulting empirical p-value (0/300, $p \approx 0.0033$ for both targets).

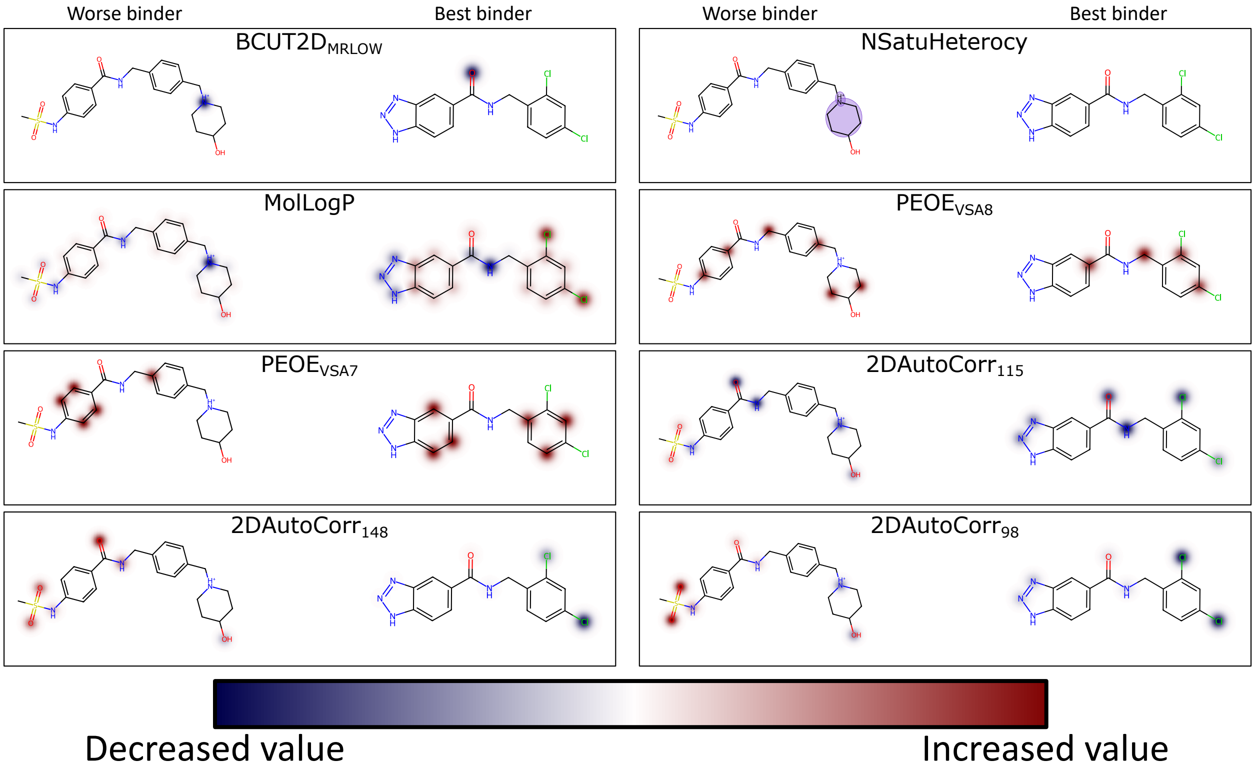


Figure S13: **Analysis of molecular-descriptor contributions to affinity in the QSAR model** Atom-level mapping of four representative descriptors from Figure S7 Panel C onto the best (253h) and worst (107t) binders in the dataset (Left panels: Increasing the descriptor improves affinity. Right panels: Increasing the descriptor hinders affinity). The color scale indicates atomic contributions toward increasing or decreasing descriptor values. Contributions for the 2DAutoCorr and BCUT2D descriptors were estimated by atom-wise carbon substitution, whereas PEOE$_{VSAx}$ contributions were obtained from their analytical atomic decompositions. The latter are non-negative by construction and therefore populate only the positive half of the color scale.

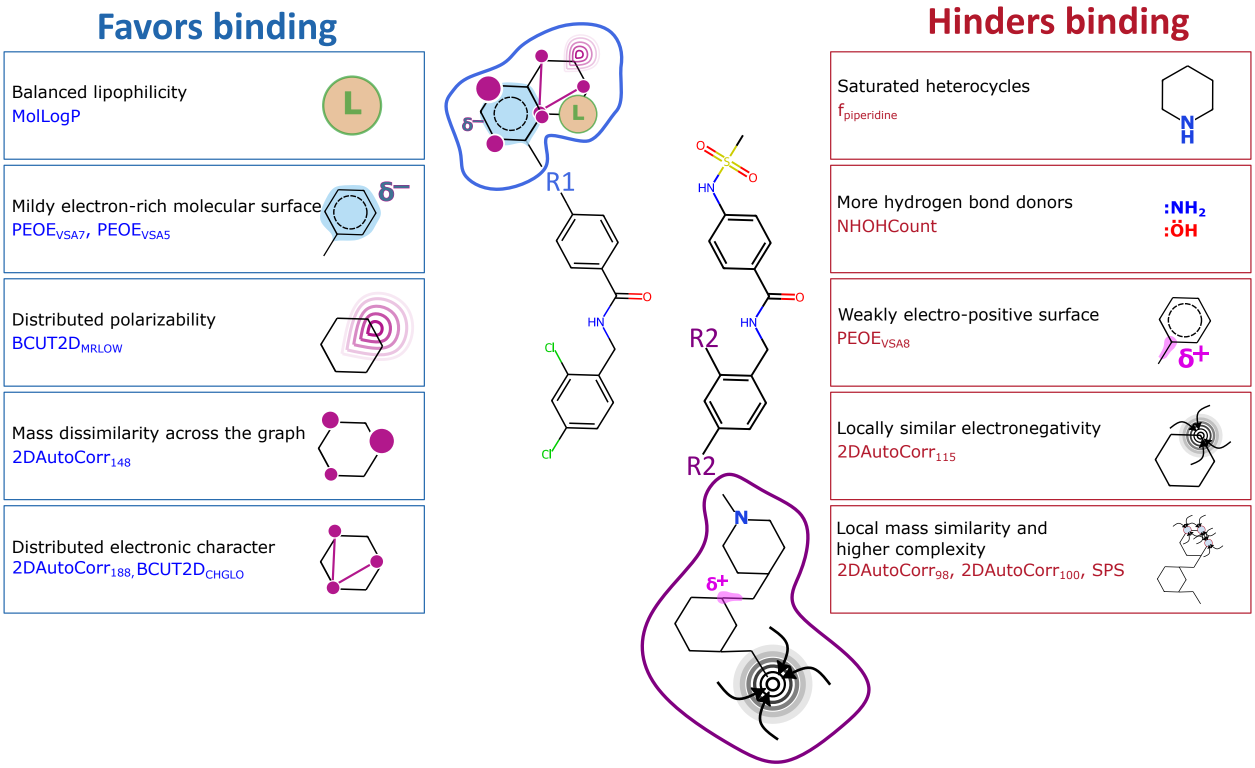


Figure S14: **Summary of molecular-descriptor contributions to affinity in the QSAR model**. Representative scheme of consensus upper (R1) and lower (R2) Q6F derivatives, based on the molecular descriptors contributing to the QSAR model. The annotations provide an overall chemical interpretation of how each descriptor may improve or hinder binding affinity.

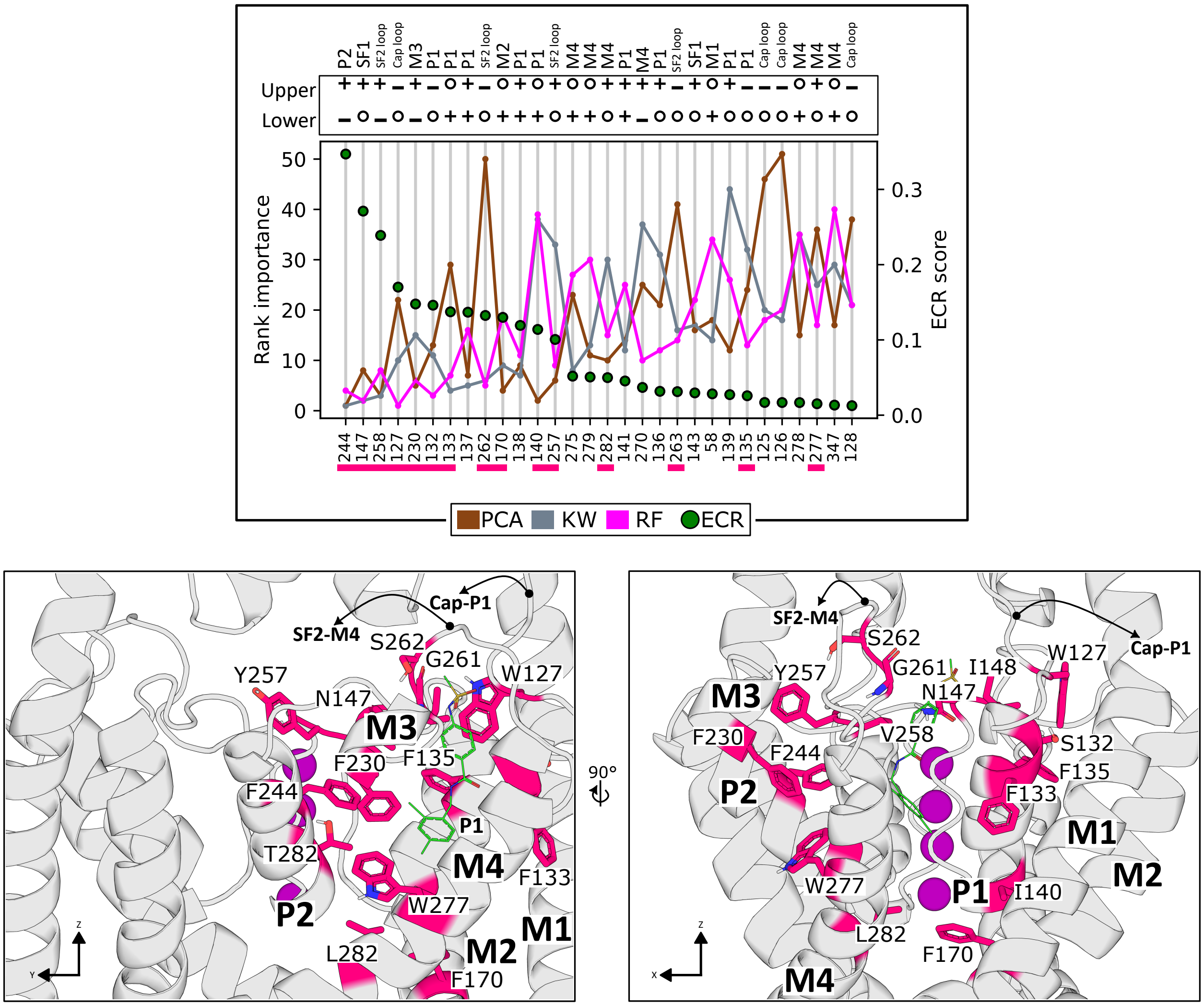


Figure S15: **Analysis of interaction patterns between Q6F derivatives and the channel. Upper panel**: Ranking of residue importance based on the interaction-pattern matrix shown in **Figure 9**. Principal component analysis (PCA) identifies the directions capturing the largest variance in the interaction patterns, and residues are ranked according to their contributions to the principal components. The Kruskal-Wallis (KW) test evaluates whether residue-level contact changes differ significantly among the three affinity groups (high, medium, and low affinity), with multiple-testing-corrected $p$-values used to rank residue significance. A random forest (RF) classifier assigns each ligand to one of the three affinity classes using differences in interaction patterns as input features, and the resulting feature-importance scores identify the residues contributing most strongly to the classification. The rankings obtained from PCA, KW, and RF were subsequently combined using exponential consensus ranking (ECR). Symbols shown above the residue rankings indicate whether upper- or lower-ring substitution leads to an increase (+), decrease (−), or no appreciable change (○) in the number of contacts relative to Q6F. Labels indicate the structural region of the channel to which each residue belongs. The magenta rectangle highlights a subset of high-ranked residues according to the ECR analysis. **Lower panel**: Structural representation of the modulator pocket with key residues highlighted. Residues shown in magenta correspond to some high-ranked residues according to the ECR score, including those emphasized by the magenta rectangle in the upper panel.

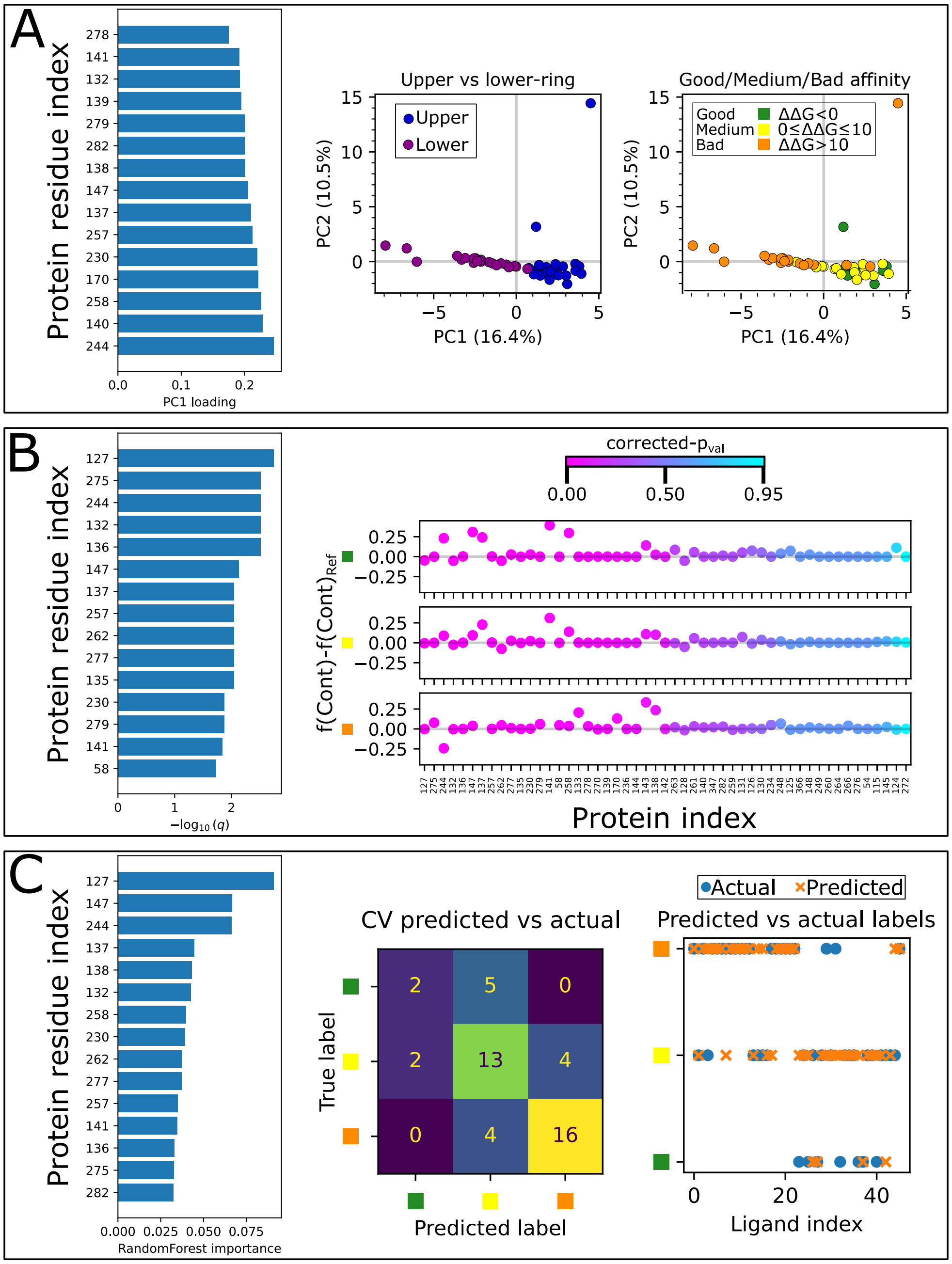


Figure S16: **Analysis of interaction patterns between Q6F derivatives and the channel. Panel A**: Principal component analysis (PCA) of the changes in ligand-residue interaction patterns. Ligands are projected onto the first two principal components and colored according to the position of the substitution (upper or lower ring; middle) or their relative-affinity class (good, medium, or poor; right). Residues with the largest contributions to PC1 are ranked according to their PC1 loadings (left). **Panel B**: Kruskal-Wallis (KW) analysis of residue-level changes in contact frequency across the three relative-affinity classes. Residues are ranked according to the significance of the differences after correction for multiple testing, expressed as $-\log_{10}(q)$ (left). Contact-frequency changes for the individual ligands are shown for the three affinity classes and colored according to the corresponding corrected $p$-value (right). **Panel C**: Random forest (RF) classification of ligands into good-, medium-, and poor-affinity classes using residue-level interaction changes as input features. Residues are ranked according to their RF impurity-based feature importance (left). Cross-validated classification performance is summarized by the predicted-versus-actual confusion matrix (middle), while the predicted and actual affinity classes for individual ligands are shown on the right.

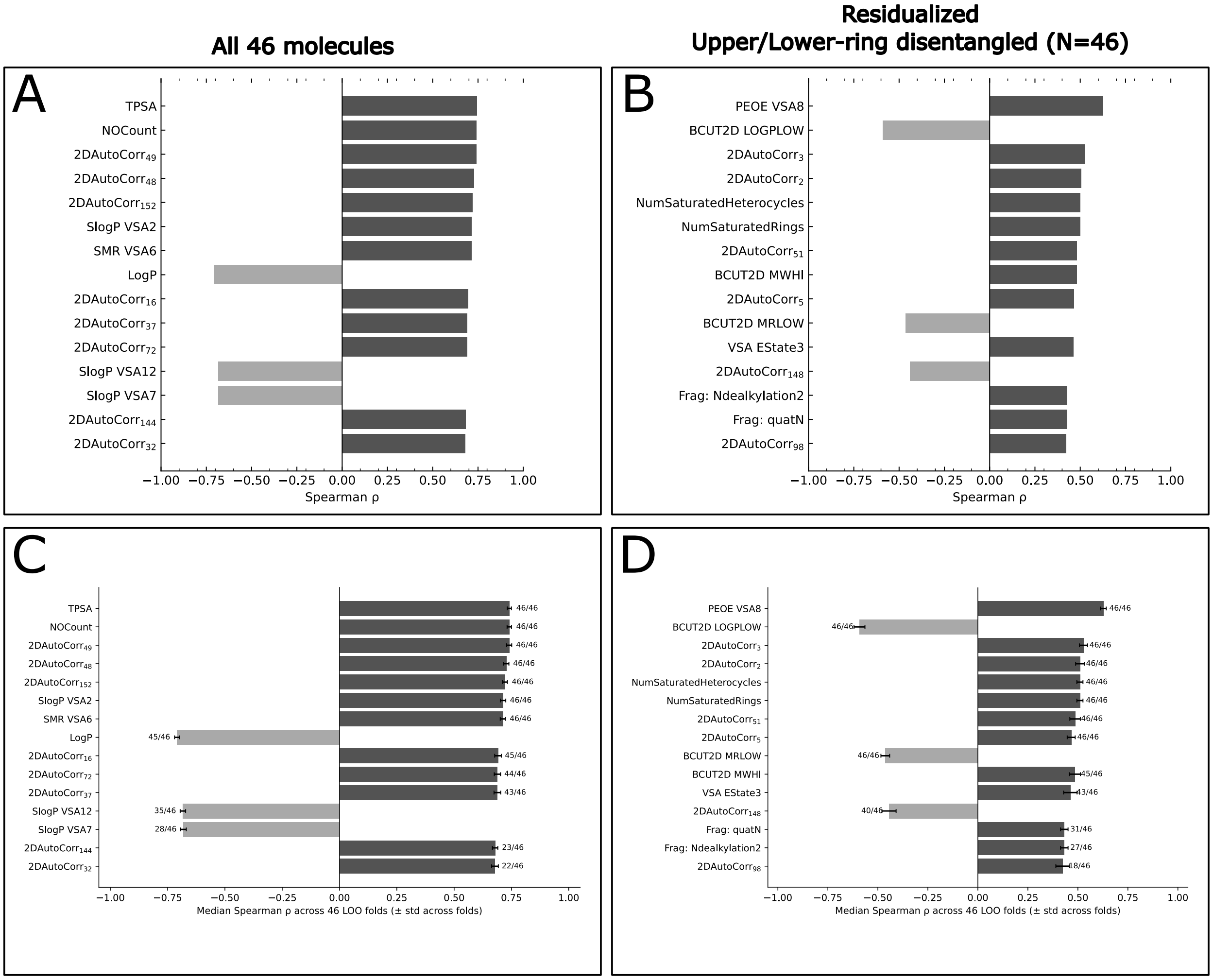


Figure S17: **Univariate descriptor correlations with $\Delta\Delta G$, before and after residualizing for Upper/Lower-ring series identity**. **Panel A**. Top-15 descriptors by Spearman $\rho$ against the original (pooled) $\Delta\Delta G$, all 46 molecules. **Panel B**. Top-15 descriptors by Spearman $\rho$ against the region-residualized $\Delta\Delta G$ (Upper/Lower-ring group mean subtracted; N=46). **Panel C**. Leave-one-molecule-out stability of the ranking in Panel A: median Spearman $\rho \pm$ SD across the 46 leave-one-out folds; the fraction of folds in which each descriptor remained in the top-15 is given alongside each bar. **Panel D**. Same leave-one-molecule-out stability analysis applied to the residualized ranking in Panel B.

Table S1: **Q6F derivatives displaying the most favorable and unfavorable relative binding affinities.** Representative extremes of the calculated relative binding free-energy differences ($\Delta\Delta G$) are shown separately for favorable and unfavorable binding. Upper- and lower-ring substitutions are denoted by $h$ and $t$, respectively. Green and orange highlighting identify favorable- and unfavorable-affinity extremes, respectively, **within the Upper/Lower groups**. The same color coding is used in Tables S2 and S3. Within each group, entries are ordered from the most favorable (most negative or closest to zero) to the least favorable $\Delta\Delta G$.

| Code | Modified ring | Substituent | Relative affinity |
|---|---|---|---|
| *Most favorable relative affinities* | | | |
| 253h | Upper | Triazole-fused indole | $-9.36 \pm 0.88$ kJ mol$^{-1}$ |
| 28h | Upper | Dichloro-hydroxyphenyl | $-8.14 \pm 1.55$ kJ mol$^{-1}$ |
| 70h | Upper | Methyl-phenyl-sulfonamide | $-2.56 \pm 1.36$ kJ mol$^{-1}$ |
| 196h | Upper | Oxyphenyl-thiazole | $-2.52 \pm 1.60$ kJ mol$^{-1}$ |
| 55t | Lower | Trifluoro-oxy | $1.17 \pm 0.87$ kJ mol$^{-1}$ |
| 28t | Lower | Difluoro | $1.31 \pm 0.89$ kJ mol$^{-1}$ |
| 106t | Lower | Trimethyl-ethyl | $5.65 \pm 1.21$ kJ mol$^{-1}$ |
| 21t | Lower | Methyl | $7.07 \pm 1.12$ kJ mol$^{-1}$ |
| *Most unfavorable relative affinities* | | | |
| 51h | Upper | Fluoride | $9.32 \pm 0.69$ kJ mol$^{-1}$ |
| 254h | Upper | Carboxylate-sulfonamide | $13.60 \pm 1.29$ kJ mol$^{-1}$ |
| 252h | Upper | Phthalimide-methyloxolane | $20.05 \pm 1.15$ kJ mol$^{-1}$ |
| 251h | Upper | Morpholine-sulfonamide-methoxy | $23.57 \pm 1.82$ kJ mol$^{-1}$ |
| 105t | Lower | Acetyl-piperazine | $55.76 \pm 2.21$ kJ mol$^{-1}$ |
| 108t | Lower | Dimethyl-piperidine | $101.95 \pm 9.41$ kJ mol$^{-1}$ |
| 52t | Lower | Piperidine | $106.19 \pm 4.63$ kJ mol$^{-1}$ |
| 107t | Lower | Hydroxy-piperidine | $121.12 \pm 10.17$ kJ mol$^{-1}$ |

Table S2: **Q6F derivatives displaying extreme desolvation and hydrophilic-hydrophobic-contact differences behavior.** Compounds highlighted in the equilibrium-ensemble analysis as displaying relatively high or low contact-fraction difference, or a high desolvation difference are summarized. Desolvation refers to the relative loss of ligand hydration waters upon transfer from solution to the TREK1 modulator pocket. Dashes indicate that a compound was not specifically identified as an extreme for the corresponding observable. Compound codes highlighted in green or orange correspond to favorable- and unfavorable-affinity extremes, respectively, as defined in Table S1. Entries are ordered by decreasing magnitude within each group, from the most extreme to progressively less extreme values.

| **Code** | **Ring** | **Substituent** | **Desolvation** | **Contacts difference** |
|---|---|---|---|---|
| *Upper-ring substitutions* | | | | |
| 252h | Upper | Phthalimide-methyloxolane | $28.55 \pm 0.64$ | — |
| 28h | Upper | Dichloro-hydroxyphenyl | $28.31 \pm 0.22$ | — |
| 147h | Upper | Difluoro-diazofurane | $28.19 \pm 0.31$ | — |
| 66h | Upper | Naphthalene | $24.26 \pm 0.22$ | — |
| 51h | Upper | Fluoride | $23.10 \pm 0.19$ | — |
| 59h | Upper | Methyl | $22.80 \pm 0.27$ | $0.252 \pm 0.006$ |
| 135h | Upper | Sulfonamide-methoxy | — | $0.271 \pm 0.009$ |
| 48h | Upper | Fluoro-methoxy | — | $0.271 \pm 0.006$ |
| 107h | Upper | Methyl-fluoride | — | $0.253 \pm 0.006$ |
| 59h | Upper | Methyl | $22.80 \pm 0.27$ | $0.252 \pm 0.006$ |
| *Lower-ring substitutions* | | | | |
| 104t | Lower | Trifluoro-amide | $28.96 \pm 0.41$ | — |
| 105t | Lower | Acetyl-piperazine | $26.20 \pm 0.49$ | — |
| 61t | Lower | Trifluoro-methoxy | $25.43 \pm 0.30$ | — |
| 106t | Lower | Trimethyl-ethyl | $19.76 \pm 0.32$ | — |
| 9t | Lower | Methoxy | $19.74 \pm 0.40$ | — |
| 52t | Lower | Piperidine | $18.63 \pm 0.53$ | — |
| 21t | Lower | Methyl | $17.69 \pm 0.18$ | $0.182 \pm 0.004$ |
| 28t | Lower | Difluoro | — | $0.228 \pm 0.011$ |
| 1t | Lower | Fluoro | — | $0.193 \pm 0.009$ |
| 21t | Lower | Methyl | $17.69 \pm 0.18$ | $0.182 \pm 0.004$ |
| 13t | Lower | Trifluoro | — | $0.180 \pm 0.023$ |

Table S3: **Q6F derivatives displaying extreme conformational-entropy penalties upon binding.** Compounds identified at the high or low extremes of the torsional-entropy proxy are summarized. Compound codes highlighted in green or orange correspond to favorable- and unfavorable-affinity extremes, respectively, as defined in Table S1. Entries are ordered by decreasing magnitude within each group, from the most extreme to progressively less extreme values.

| **Code** | **Ring** | **Substituent** | **Relative entropy penalty [J mol$^{-1}$ K$^{-1}$]** |
|---|---|---|---|
| *Low conformational-entropy penalty* | | | |
| 253h | Upper | Triazole-fused indole | -27.11 $\pm$ 0.64 |
| 59h | Upper | Methyl | -23.27 $\pm$ 0.56 |
| 252h | Upper | Phthalimide-methyloxolane | -18.47 $\pm$ 0.59 |
| 3t | Lower | Benzodioxole | -18.80 $\pm$ 0.48 |
| 52t | Lower | Piperidine | -18.72 $\pm$ 0.60 |
| 66t | Lower | Ethoxy-pyridine | -14.58 $\pm$ 0.57 |
| *High conformational-entropy penalty* | | | |
| 237h | Upper | fluor-methoxy | -32.57 $\pm$ 0.54 |
| 219h | Upper | di-methyl-oxazole | -32.46 $\pm$ 0.52 |
| 107h | Upper | Methyl-fluoride | -32.30 $\pm$ 0.56 |
| 189h | Upper | sulfo-piperidine | -31.97 $\pm$ 0.53 |
| 77t | Lower | Pyrazole | -32.25 $\pm$ 0.55 |
| 35t | Lower | Imidazole | -31.77 $\pm$ 0.55 |
| 55t | Lower | Trifluoro-oxy | -31.10 $\pm$ 0.53 |

Table S4: **Molecular descriptors contributing to the consensus QSAR model.** Descriptors are grouped according to the sign of their mean ElasticNet coefficient across the LOOCV models. Negative coefficients are associated with lower $\Delta\Delta G$ values and therefore improved relative affinity, whereas positive coefficients are associated with higher $\Delta\Delta G$ values and reduced relative affinity. The final column indicates descriptors for which an atom-level interpretation is discussed in Figure S9.

| **Descriptor** | **Descriptor definition** | **Chemical interpretation in this series** | **Atom-level example** |
|---|---|---|---|
| *Descriptors associated with improved affinity* | | | |
| $BCUT2D_{MRLOW}$ | Lowest eigenvalue of the BCUT connectivity matrix weighted by molar refractivity | Distribution of polarizability and atomic volume across the molecular graph | Yes |
| MolLogP | Calculated octanol/water partition coefficient | Greater lipophilicity within the sampled chemical space is associated with improved affinity | Yes |
| QED | Composite quantitative estimate of drug-likeness | Drug-like combinations of physicochemical properties are associated with improved affinity | No |
| $PEOE_{VSA7}$ | Van der Waals surface area from atoms with Gasteiger partial charges between $-0.05$ and 0.00 | Weakly polarized, nearly neutral surface, including non-polar aliphatic and unsubstituted aromatic carbons, is favored | Yes |
| $PEOE_{VSA5}$ | Van der Waals surface area from atoms with Gasteiger partial charges between $-0.15$ and $-0.10$ | Moderately electron-rich molecular surface is associated with improved affinity | No |
| $2DAutoCorr_{148}$ | Geary autocorrelation of relative atomic mass for atoms separated by four bonds | Greater mass heterogeneity at this topological separation is associated with improved affinity | Yes |

Continued on next page

**Table S4 continued**

| Descriptor | Descriptor definition | Chemical interpretation in this series | Atom-level example |
|---|---|---|---|
| $EState_{VSA3}$ | Van der Waals surface area from atoms with electrotopological-state values between 0.29 and 0.72 | Represents relatively weakly polarized atomic environments | No |
| $BCUT2D_{CHGLO}$ | Lowest eigenvalue of the BCUT matrix weighted by Gasteiger partial charges | Distribution of partial-charge information throughout the molecular graph | No |
| $2DAutoCorr_{188}$ | Geary autocorrelation of electrotopological state for atoms separated by four bonds | Greater electronic heterogeneity across the molecular graph is associated with improved affinity | No |
| $EState_{VSA7}$ | Van der Waals surface area from atoms with electrotopological-state values between 1.81 and 2.05 | Surface contributed by electronically or topologically differentiated atomic environments is favored | No |
| *Descriptors associated with reduced affinity* | | | |
| $fr_{Ndealkylation2}$ | SMARTS-based fragment descriptor identifying tertiary alicyclic amine motifs | Nitrogen-containing saturated heterocycles are associated with poorer affinity[a] | No |
| $fr_{piperidine}$ | SMARTS-based count of piperidine motifs | Piperidine-containing derivatives are associated with poorer affinity[a] | No |
| $N_{SatuHeterocy}$ | Number of saturated heterocyclic rings | A larger number of saturated heterocycles is associated with reduced affinity | Yes |
| $PEOE_{VSA8}$ | Van der Waals surface area from atoms with Gasteiger partial charges between 0.00 and +0.05 | Greater weakly electropositive surface is associated with reduced affinity | Yes |

Continued on next page

**Table S4 continued**

| Descriptor | Descriptor definition | Chemical interpretation in this series | Atom-level example |
|---|---|---|---|
| 2DAutoCorr$_{115}$ | Moran autocorrelation of atomic electronegativity for atoms separated by four bonds | Topological clustering of atoms with similar electronegativity is associated with reduced affinity | Yes |
| NHOHCount | Number of NH and OH groups | More potential hydrogen-bond donors are associated with poorer affinity, possibly reflecting an unfavorable polarity/desolvation balance | No |
| EState$_{VSA5}$ | Van der Waals surface area from atoms with electrotopological-state values between approximately 1.17 and 1.54 | Represents environments commonly associated with carbon atoms influenced by nearby electronegative atoms | No |
| 2DAutoCorr$_{98}$ | Moran autocorrelation of relative atomic mass for atoms separated by three bonds | Greater short-range similarity in atomic mass is associated with reduced affinity | Yes |
| 2DAutoCorr$_{100}$ | Moran autocorrelation of relative atomic mass for atoms separated by five bonds | Longer-range atomic-mass correlation is associated with reduced affinity | No |
| SPS | Spatial complexity based on sp$^3$ character, hybridization, stereochemistry, ring membership, and connectivity | Greater structural and topological complexity is associated with reduced affinity | No |

[a] The large coefficients of fr$_{Ndealkylation2}$ and fr$_{piperidine}$ should be interpreted cautiously because these fragments occur primarily in a small number of poorly performing derivatives and may therefore reflect extreme compounds rather than a broadly transferable chemical trend.

### Supplementary Methods: Total contact fraction analysis

Further information can be extracted from the contact matrix difference using unsupervised, statistical, and supervised analyses, as follows:

- **Principal Component Analysis (PCA)**: In this case, the contact matrix $\mathbf{X}$ is standardized by subtracting the mean and dividing by the standard deviation of each feature. Given the standardized matrix $\mathbf{X} \in \mathbb{R}^{n \times p}$ (with zero mean and unit variance for each feature), the covariance matrix of the standardized data is computed as

$$\mathbf{C} = \frac{1}{n-1}\mathbf{X}^{\top}\mathbf{X}. \tag{1}$$

Since the variables are standardized, this matrix is equivalent to the correlation matrix. PCA is then carried out through the eigendecomposition of $\mathbf{C}$:

$$\mathbf{C}\mathbf{v}_k = \lambda_k \mathbf{v}_k, \tag{2}$$

where $\mathbf{v} * k$ are the eigenvectors and $\lambda * k$ the corresponding eigenvalues, ordered such that $\lambda_1 \geq \lambda_2 \geq \cdots \geq \lambda_p$. The eigenvectors define the principal components, i.e., the orthogonal directions of maximal variance. The projection of the data onto these directions provides the scores:

$$\mathbf{Z} = \mathbf{X}\mathbf{V}, \tag{3}$$

where $\mathbf{V} = [\mathbf{v}_1, \mathbf{v}_2, \ldots, \mathbf{v}_p]$ is the matrix of eigenvectors, and $\mathbf{Z}$ contains the representation of the data in principal-component space. The coefficients defining the contribution of each original variable to a principal component are referred to as *loadings*. Thus, the loading of a given residue-level contact feature indicates the magnitude and direction of its contribution to the variance captured by that component, with larger absolute loading values identifying features that contribute more strongly to the corresponding principal component.

- **Kruskal-Wallis test (KW)**: The Kruskal-Wallis test[48] is a non-parametric rank-based test used to compare two or more independent groups, allowing for unequal sample sizes. Here, for each residue, the contact differences were partitioned into three groups according to ligand affinity: good ($\Delta\Delta G < 0$ $kJ/mol$), medium ($0$ $kJ/mol \leq \Delta\Delta G \leq 10$ $kJ/mol$), and bad ($\Delta\Delta G > 10$ $kJ/mol$). The test was then used to assess whether the groups are statistically indistinguishable or whether at least one group tends to exhibit systematically different values. More precisely, the null hypothesis is that the group samples come from the same distribution; when group distributions have similar shape and dispersion, the test is commonly interpreted as probing differences in central tendency. The resulting $p$-values were corrected for multiple testing using a false discovery rate (FDR) procedure.[49]

- **Random forest (RF)**: A random-forest classifier[50] was trained to predict ligand classes (good, medium, bad, as in the KW test) from the residue-level contact differences. Briefly, a nested cross-validation scheme was employed using stratified $K$-fold splits ($n = 3$): the inner loop was used for hyperparameter optimization, and the outer loop for unbiased performance estimation. Hyperparameters were tuned using randomized search over a predefined parameter space, optimizing balanced accuracy in order to account for class imbalance. Model performance was then assessed using the nested cross-validated balanced accuracy (mean $\pm$ standard deviation), cross-validated predictions, the classification report, and the confusion matrix.

  Finally, the model was retrained on the full dataset using the optimal hyperparameters. The number of trees was increased in order to stabilize the feature-importance estimates. Residue importance was quantified using impurity-based (Gini) feature importance, ranking residues according to their contribution to the classification.

For each of these approaches, an importance measure was extracted; namely, the absolute value of the loadings of each residue contact difference on the first principal component in PCA, the statistical evidence associated with each residue in the KW test, and the impurity-based feature importance from the RF classifier. In the KW case, because smaller corrected $p$-values indicate stronger evidence, the statistical importance was quantified as $-\log_{10}(q)$, where $q$ denotes the FDR-corrected $p$-value. These importance measures were then normalized internally by subtracting the minimum value and dividing by the corresponding range.

To obtain a consensus across methods, we applied an Exponential Consensus Ranking (ECR).[51] This method combines multiple independent rankings into a single consensus score while emphasizing top-ranked elements. Instead of weighting all rank positions equally, ECR assigns exponentially decaying weights so that higher-ranked items contribute much more strongly than lower-ranked ones.

For a given element $i$ and ranking method $j$, an exponential score is assigned based on its rank $r_i^{(j)}$. The final consensus score is then obtained by summing these contributions across all ranking methods. This ensures that residues consistently appearing among the top ranks across different methods are prioritized, whereas low-ranked residues contribute only weakly.

The parameter $\sigma$ controls the rate of exponential decay and therefore how strongly top-ranked elements are favored in the consensus (in this case, $\sigma$=30). The ECR score is defined as

$$p\left(r_i^{(j)}\right) = \frac{1}{\sigma}\exp\left(-\frac{r_i^{(j)}}{\sigma}\right), \tag{4}$$

and the final consensus score for residue $i$ is

$$P(i) = \sum_j p\left(r_i^{(j)}\right) = \frac{1}{\sigma}\sum_j \exp\left(-\frac{r_i^{(j)}}{\sigma}\right). \tag{5}$$

### Supplementary Methods: Penalized linear regression model (QSAR) construction

To understand the relationship between certain physicochemical properties of the ligands and their calculated binding free energies, molecular descriptors were combined with penalized linear regression and univariate correlation analyses. A total of 414 physicochemical and topological molecular descriptors were calculated for the 46 molecules using the RDKit package.[42] The molecules belonged to two structurally distinct derivative series: Upper and lower-ring substitutions. Descriptors with zero variance across the complete dataset were removed before subsequent analyses.

Because the number of molecular descriptors was larger than the number of molecules, an additional decorrelation step was applied before modeling. Pairwise correlations between descriptors were converted into a distance matrix according to $1 - |r|$, and descriptors were hierarchically clustered using average linkage. Descriptors belonging to the same cluster at an absolute correlation threshold of $|r| > 0.95$ were considered redundant, and the descriptor with the largest variance within each cluster was retained as its representative. The final decorrelated descriptor matrix was used for the ElasticNet analysis. Univariate correlation analyses were instead performed using the complete set of non-constant descriptors.

Both upper and lower-ring derivative series display, on average, a pronounced difference in their relative binding-free energies. Consequently, a model trained directly on the pooled binding free energies could potentially capture structural features distinguishing the two series rather than relationships between molecular properties and affinity within each series.

Because the Upper- and Lower-ring series differ systematically in mean binding affinity (See Results), a residualized binding free-energy target was constructed. For each molecule, the mean binding free energy of the corresponding Upper or Lower-ring series was subtracted from its calculated value,

$$\Delta\Delta G_i^{\text{res}} = \Delta\Delta G_i - \langle \Delta\Delta G \rangle_{r(i)}\,, \tag{6}$$

where $r(i)$ denotes the Upper or Lower-ring series to which molecule $i$ belongs. The residualized affinity represents variation in binding affinity relative to the mean of the corresponding derivative series. ElasticNet models were fitted for both the original and residualized targets to assess how much of the predictive signal was retained after removal of the between-series contribution.

Predictive performance was evaluated using a nested leave-one-out cross-validation (LOOCV) procedure. The outer cross-validation consisted of 46 folds, such that one molecule was excluded in each fold and the remaining 45 molecules were used for model fit. Descriptor standardization was performed independently within each outer training fold using the mean and standard deviation of the training data only, and the same transformation was subsequently applied to the held-out molecule. This procedure prevented information from the test molecule from contributing to descriptor scaling.

Within each outer training fold, the ElasticNet regularization parameters were selected using an inner five-fold cross-validation procedure with shuffled folds. Seven values of the L1/L2 mixing parameter were explored ($\rho$ = 0.10, 0.30, 0.50, 0.70, 0.85, 0.95, and 1.00), together with 50 candidate values of the regularization strength $\alpha$. The fitted model was then used to predict the binding free energy of the molecule excluded from the outer fold, resulting in one strictly out-of-fold prediction for each of the 46 molecules.

The quality of the cross-validated predictions was quantified using the leave-one-out cross-validated coefficient of determination ($Q^2_{\text{CVLOO}}$), the mean absolute error (MAE), and the Spearman rank correlation coefficient ($\rho$) between the observed and out-of-fold predicted values. These quantities were calculated as

$$Q^2_{\text{LOO}} = 1 - \frac{\sum_{i=1}^{n} (y_i - \hat{y}_i)^2}{\sum_{i=1}^{n} (y_i - \bar{y})^2}\,, \tag{7}$$

where $y_i$ is the observed binding free energy of molecule $i$, $\hat{y}_i$ is its prediction calculated when that molecule was excluded from model fitting, and $\bar{y}$ is the mean observed binding free energy across the complete dataset. The mean absolute error was calculated as

$$\text{MAE} = \frac{1}{n}\sum_{i=1}^{n} |y_i - \hat{y}_i|\,, \tag{8}$$

providing the mean absolute deviation between the observed and out-of-fold predicted binding free energies. The Spearman rank correlation coefficient was calculated between the observed and predicted values after conversion of both variables to ranks. It can be expressed as

$$\rho = \frac{\sum_{i=1}^{n} \left(R_i - \bar{R}\right)\left(\hat{R}_i - \bar{\hat{R}}\right)}{\sqrt{\sum_{i=1}^{n} \left(R_i - \bar{R}\right)^2 \sum_{i=1}^{n} \left(\hat{R}_i - \bar{\hat{R}}\right)^2}}, \tag{9}$$

where $R_i$ and $\hat{R}_i$ are the ranks of the observed and predicted binding free energies, respectively. These metrics were evaluated independently for models fitted from the original and residualized binding free energies.

To determine whether the observed predictive performance could arise by chance in the high-dimensional descriptor space, a permutation analysis was additionally performed for both targets. Binding free-energy values were randomly permuted among the molecules and the nested LOOCV procedure was repeated to generate an empirical null distribution of the cross-validated $Q^2$ (**Figure S17**). The permutation $p$-value was estimated as

$$p_{\text{perm}} = \frac{1 + \sum_{k=1}^{N_{\text{perm}}} I\left(Q^2_{k,\text{null}} \geq Q^2_{\text{obs}}\right)}{N_{\text{perm}} + 1}. \tag{10}$$

A total of 300 permutations were used in the final calculations. For computational efficiency, the permutation analysis used the complete decorrelated descriptor set but a reduced inner hyperparameter search consisting of two L1 ratios (0.5 and 1.0), 10 candidate values of $\alpha$, and two-fold inner cross-validation. This approximation was applied only to construction of the permutation null distribution; the models used to estimate the reported predictive performance employed the full hyperparameter search described above.

Because a single exhaustive LOOCV calculation produces only one prediction for each molecule, the sensitivity of individual predictions to the choice of the inner cross-validation split was evaluated separately. The complete nested LOOCV procedure was repeated 10 times using different random seeds for the shuffled inner five-fold cross-validation. The outer leave-one-out partition remained unchanged in all repetitions. For each molecule, the mean and standard deviation of its 10 out-of-fold predictions were calculated. The resulting standard deviation was interpreted as a measure of the uncertainty associated with the inner cross-validation and hyperparameter-selection procedure, rather than as a formal statistical confidence interval.

The stability of the descriptors selected by the ElasticNet model was also evaluated across the 46 outer LOOCV folds. For each fold, the standardized regression coefficients of the fitted model were retained. For each descriptor, the mean and standard deviation of the coefficient across all 46 folds were calculated, with folds in which the descriptor was not selected included as coefficients equal to zero. Descriptor selection frequency was defined as the fraction of LOOCV folds in which the absolute coefficient was greater than $10^{-8}$. This analysis therefore quantified both the magnitude and reproducibility of the contribution of individual descriptors across independent perturbations of the training dataset, without relying on a separate bootstrap procedure.

As a complementary model-independent analysis, Spearman rank correlations were calculated between each individual molecular descriptor and the binding free-energy target. Correlations were evaluated both for the original binding free energies and for the residualized values using all 46 molecules. For each descriptor, the corresponding two-sided $p$-value was calculated and corrected for multiple testing using the Benjamini-Hochberg false-discovery rate procedure.[49] Descriptors were ranked according to the absolute magnitude of their Spearman correlation coefficient.

The robustness of this univariate descriptor ranking to individual molecules was further examined using an exhaustive leave-one-molecule-out stability analysis. In each of the 46 folds, one molecule was removed and, for the residualized analysis, the Upper and Lower-ring series means were recalculated using only the remaining 45 molecules. Spearman correlations were then recomputed between all descriptors and the corresponding residualized binding free energies, and the 15 descriptors with the largest absolute correlation coefficients were recorded. The selection fraction of a descriptor was defined as the ratio of the 46 folds in which it appeared among these top 15 descriptors. In addition, the median Spearman coefficient and its standard deviation across folds were calculated. Descriptors consistently recovered across the leave-one-out folds were considered robust univariate correlates of binding affinity.

The penalized linear regression model was based on the ElasticNet approach,[52] which combines L1 and L2 regularization. The L1 component promotes sparse solutions by shrinking some regression coefficients exactly to zero, whereas the L2 component reduces the magnitude of correlated coefficients and stabilizes their estimation. The ElasticNet estimator was defined as

$$\hat{\boldsymbol{\beta}} = \underset{\boldsymbol{\beta} \in \mathbb{R}^p}{\arg\min} \left\{ \frac{1}{2n} \left\| \mathbf{y} - \mathbf{X}\boldsymbol{\beta} \right\|_2^2 + \alpha\rho \left\| \boldsymbol{\beta} \right\|_1 + \frac{1}{2}\alpha(1-\rho) \left\| \boldsymbol{\beta} \right\|_2^2 \right\}, \tag{11}$$

where $\mathbf{y}$ is the vector of binding free-energy values, $\mathbf{X}$ is the standardized molecular-descriptor matrix, $\boldsymbol{\beta}$ is the vector of regression coefficients, $\alpha$ controls the total strength of regularization, and $\rho$ controls the relative contributions of the L1 and L2 penalties. The limiting case $\rho = 1$ corresponds to Lasso regression, whereas $\rho = 0$ corresponds to Ridge regression. In the present analysis, both $\alpha$ and $\rho$ were optimized exclusively within the training data of each outer LOOCV fold.

We further estimated the contribution of individual atoms to selected molecular descriptors. For descriptors that are analytically defined as sums of atomic contributions, these values were mapped directly onto the molecular structure. For $PEOE_{VSAx}$, atomic contributions correspond to each atom's Labute approximate surface-area (ASA) increment, restricted to atoms whose Gasteiger (PEOE) partial charge falls within the charge interval defining the corresponding bin. For MolLogP, atomic contributions correspond to the additive Crippen atom-type increments to the octanol/water partition coefficient,[53] which sum exactly to the whole-molecule value.

For descriptors without an exact atomic decomposition, including the 2DAutoCorr and BCUT2D descriptors; atomic contributions were estimated using an atom-wise perturbation approach. Each atom was replaced in turn by a carbon atom while keeping the molecular topology fixed, after which the descriptor was recalculated. The resulting change in descriptor value was used as a local measure of the contribution of the chemical identity and environment of that atom. In the case of $BCUT2D_{MRLOW}$, which corresponds to an extreme eigenvalue of a modified adjacency matrix whose diagonal encodes atomic molar refractivity and whose off-diagonal elements encode bond connectivity, this perturbation captures both the direct change in the atomic property and the indirect influence of the atom on the overall matrix spectrum.

### Supplementary Methods: Parameter refinement methodology

For the two molecules with experimentally measured values (Q6Fa and Q5F), a fragment-based parameter optimization approach was implemented. Fragments corresponding to the N-cyclohexylsulfonamide moiety of Q6Fa and the N-methyl-thiophene-2-carboxamide moiety of Q5F were optimized using the standard CHARMM General Force Field[24–26] (CGenFF) parameterization workflow, as implemented in the Force Field Toolkit[27,54,55] (ffTK) plugin of VMD.

Initial bonded and non-bonded parameters for the two molecules were derived by analogy using the CGenFF program, as implemented in the CHARMM-GUI web server, together with their corresponding CGenFF penalty scores. Because the highest penalties were localized in specific regions of the molecules (the N-$SO_2$ linker in Q6Fa and the amide-thiophene linker in Q5F, with penalties of 10.9/35.1 and 4.9/129 for charges and torsions, respectively), reoptimization was restricted to these fragments rather than applied to the complete molecules. All quantum-mechanical (QM) calculations described below were carried out on the isolated fragments, capped as neutral closed-shell singlets, using Gaussian 09.[56]

Fragment geometries were optimized at the MP2/6-31G(d) level (`Opt=Redundant`, `SCF=Tight`). Analytic Hessians were calculated at the same level (MP2/6-31G(d), `Freq`, tight SCF) and used within the ffTK BondedOpt module to obtain initial equilibrium values and force constants for bonds and angles centered on the flagged atoms. These parameters were then refined by simplex minimization against the QM Hessian.

Partial atomic charges were optimized using the water-interaction ChargeOpt module of ffTK. For each polar donor or acceptor site in the fragment, a fragment-water complex was set up and its geometry optimized at the HF/6-31G(d) level (`Opt=Z-matrix`), with an isolated water molecule treated at the RHF/6-31G(d) level as reference. Trial charges on atoms neighboring each flagged site were iteratively adjusted by downhill-simplex minimization of an objective function combining (i) the deviation between MM and QM interaction energies and heavy-atom-water distances for each donor/acceptor geometry and (ii) the deviation of the MM molecular dipole moment from the QM dipole moment calculated at the optimized fragment geometry. Charges outside the reparameterized substructure were retained at their original CGenFF analogy values, and charge neutrality of the fragment was enforced.

Dihedral parameters spanning the reparameterized region (four dihedrals for Q6Fa and three for Q5F) were obtained from relaxed potential-energy surface scans at the MP2/6-31G(d) level. Scans were performed bidirectionally from the optimized geometry in $15°$ increments over a $\pm 90°$ range using `Opt=ModRedundant`. Torsional Fourier coefficients (force constant, multiplicity, and phase) were fitted to the QM energy profiles by simplex/least-squares minimization using the ffTK DihOpt module. For Q5F, whose torsional parameters carried the highest CGenFF penalty score (129), an additional refinement step using DihOptRefine was performed. In this step, the Fourier coefficients were reoptimized against the full QM torsional energy surface while keeping the updated charge and bonded parameters fixed, improving the agreement beyond the initial single-scan fit.

All optimized bonded, torsional, and charge parameters were finally combined with the unmodified CGenFF & CHARMM36m parameters describing the remainder of each ligand.

### Supplementary Discussion: Contact matrix analysis

To extract a more quantitative and interpretable picture of the relationship between contact patterns and binding affinity, we carried out three complementary analyses on the contact data (**Figure 4**): (i) a Principal Component Analysis (PCA), to identify which features (in this case, which residue contact fraction differences) contribute most to the directions of variance; (ii) a non-parametric Kruskal–Wallis (KW) test, to compare three independent groups and assess whether they derive from the same underlying distribution. The three groups are defined based on affinity (Bad binder: $\Delta\Delta G > 10\ kJ/mol$. Medium binder: $0\ kJ/mol \leq \Delta\Delta G \leq 10\ kJ/mol$. Good binder: $\Delta\Delta G < 0\ kJ/mol$). Each residue of the channel then carries an average contact fraction difference value within each group, and the test identifies which groups are statistically distinguishable, quantified by the corrected p-value; and (iii) a supervised machine-learning approach using a Random Forest (RF) classifier, which assigns each molecule (on the basis of its contact matrix) to one of the three affinity labels.

Each of these analyses provide a ranking of residues by internal "importance": the weight on the principal component in PCA, the corrected p-value in the KW test, and the feature importance in the RF classifier. After standardizing these importance scores, the three methods converge on a consistent set of residues, in agreement with our earlier qualitative analysis: residues in the upper-middle plane of the pocket (spanning the upper-middle P1 region, the M3–P2 interface, and the SF2-M4 loop), emerge as more important than, for instance, M4 residues (**Figure S10**).

### Supplementary Discussion: QSAR description

As an exercise to rationalize the contribution of the descriptors to the QSAR model, we selected four representative descriptors associated with favorable and unfavorable changes in the free-energy difference and mapped their atomic contributions onto the 2D structures of one of the best and one of the worst binders in the tested derivative set (**Figure S13**).

From the descriptors contributing to better affinity, $\mathrm{BCUT2D_{MRLOW}}$ shows that replacing the positively charged nitrogen in the piperidine ring of the worse binder, or the carbonyl oxygen in the better binder, with a carbon atom, increases the value of the descriptor (which in turn should improve affinity). This suggests that atoms with lower molar refractivity at specific topological positions in the molecular graph can hinder binding, an effect that is particularly apparent for the positively charged nitrogen in the worse binder. MolLogP shows a clearer trend: in the worse binder, the positively charged nitrogen in the piperidine ring contributes negatively to the total descriptor and therefore to a less favorable affinity. In contrast, the better binder contains both positive contributions, mainly from the two aromatic rings and chlorine substituents, and negative contributions, particularly from the NH and $NH_2$ groups in the middle and upper parts of the molecule. Together, these contributions provide a more balanced distribution of lipophilicity and, consequently, a more favorable contribution to the free energy of binding.

$\mathrm{PEOE_{VSA7}}$ further suggests that van der Waals surface area contributed by slightly electron-rich atoms, such as aromatic carbons, is associated with a higher descriptor value and then, with better binding. The better binder displays a larger contribution from these chemical environments, whereas the worse binder contains fewer of them as a consequence of the lower-ring substitution. Finally, the $\mathrm{2DAutoCorr_{148}}$ descriptor shows that replacing the $SO_2$ group and the carbonyl oxygen in the middle-upper region of the molecule with a carbon atom increases the difference between the perturbed and unperturbed molecule, this means, decreases the absolute value of the descriptor for the unperturbed molecule. Because this descriptor captures mass dissimilarity across the molecular graph at intermediate topological distances, these perturbations suggest that modifying the environment around these groups reduces the corresponding mass-distribution signal and is associated with worse affinity. In the best binder, replacing the chlorine substituent in the lower ring with carbon instead increases the absolute value of the descriptor and is associated with improved affinity. This suggests that, in this particular molecular context, the chlorine substituent contribute less favorably to the mass-dissimilarity pattern captured by the descriptor.

The analysis can likewise be extended to descriptors associated with worse affinity. $N_{\mathrm{SatuHeterocy}}$ indicates that saturated heterocycles, such as the piperidine ring present in the worse binder, are associated with reduced affinity, whereas the best binder does not display such a substitution. $PEOE_{\mathrm{VSA8}}$ suggests that van der Waals surface area contributed by weakly positive atoms is detrimental to binding. These atoms are commonly carbons adjacent to a heteroatom or to an aromatic system. In this case, the worse binder contains a larger number of such atomic environments, resulting in a larger contribution to the descriptor and, ultimately, a less favorable affinity.

The $2DAutoCorr_{115}$ descriptor captures electronegativity similarity across the molecular graph (4-bonds apart). For the worse binder, replacing the $NH_2CO$ group in the central region, the positively charged nitrogen and OH group in the piperidine ring, and the $NHSO_2$ group in the upper ring with carbon increases the value of the descriptor in the perturbed molecule, which is associated with worse affinity. A qualitatively similar pattern appears for the best binder, although to a lesser extent, with the largest effects localized around the nitrogen atoms in the upper ring, the $NH_2CO$ group in the central region, and the chlorine substituents in the lower ring. Overall, this suggests that a greater variation in electronegativity across these molecular regions is associated with improved binding.

Finally, $2DAutoCorr_{98}$ describes a related topological property, but based on atomic mass rather than electronegativity and evaluated over a shorter topological distance of three bonds apart. Here, the behavior is somewhat different. In the worse binder, the $SO_2$ group in the upper ring increases the descriptor value relative to their corresponding carbon-substituted structures. This is consistent with these groups contributing to a local mass-correlation pattern associated with worse affinity in the original molecule. In the better binder, the chlorine substituents in the lower ring make the dominant contribution: replacing them with carbon increases the absolute value of the descriptor, which is itself associated with worse affinity. Thus, in this particular molecular environment, the chlorine substituents appear to oppose the short-range mass-correlation pattern that is penalized by the model.

Overall, this comparison suggests that lower-ring substitutions can cause the molecule to lose several properties associated with favorable binding, including surface area contributed by moderately electron-rich atoms, overall lipophilicity, the topological distribution of molar refractivity, and favorable patterns of mass dissimilarity across the molecular graph. In contrast, upper-ring substitutions tend to preserve these characteristics, particularly the contribution of electron-rich aromatic environments to molecular surface area and a more balanced distribution of lipophilicity. Likewise, lower-ring substitutions involving saturated heterocycles can increase the surface area contributed by slightly positively charged atoms and promote topological clusters of atoms with similar mass or electronegativity. Both features are associated here with worse affinity. Upper-ring substitutions appear less prone to introducing these detrimental features, particularly because they preserve the aromatic nature of the lower ring.

We now turn to the analysis of the Q6F-derivatives in terms of the molecular descriptors used to define them in the regression section (**Figures S11 & S12**). The interpretation of our penalized regression model suggest that aromatic delocalized systems, that display a distributed polarizability and mild electron-rich group surface are preferred (according to $\mathrm{BCUT2D_{MRLOW}}$, and $\mathrm{PEOE_{VSA7,VSA5}}$) for the prediction of better affinity. On the same line, moderate lipophilic molecules, that can still partition on more hydrophobic phases but without being completely hydrophobic, are also desired, such as aromatic groups with polar heteroatom substitutions (this is according to MolLogP).

On the other side of the spectrum, physicochemical features from the ligand that are detrimental for affinity are represented by saturated heterocyclic rings, specially substituted by nitrogen ($\mathrm{f_{piperidine}}$, NHOHCount), and by localized, heavy-atom patches ($\mathrm{2DAutoCorr_{98,100}}$), indicating that localized bulky groups are not desired, and by a complex molecular topology (SPS). Similarly, electro-positive surface patches are also penalized ($\mathrm{PEOE_{VSA8}}$ descriptor), likely because the amphipathic modulator pocket cannot satisfy completely the screening of such charged surface.

Therefore, the physicochemical features from the ligand perspective that contribute to the affinity correspond to aromatic, moderately polar (mild electron-rich) substituents (especially in the upper ring), in a way the amphipathic character is preserved. Also, one should ensure that polarizability are distributed across the molecule graph (not localized). Features to avoid should be bulky saturated heterocyclic rings, strong localized electron-positive patches, complex molecular topologies, and small groups that cannot fulfill all the required contacts in the pocket. Our hypothesis of bulkier and hydrophobic groups in the lower ring, and polar or charged groups in the upper ring as drivers of affinity turn out to be not completely accurate, since best binders are not extreme in any descriptor. They are balanced systems that distribute polarizability across ring systems, balance hydrophobic and polar contributions through heteroatom substitution (achieving moderate lipophilicity), avoid localized charge or spatially concentrated atomic volume (steric bulk) that would clash with the binding pocket geometry, and display mild electron-rich surfaces.

### Supplementary Discussion: Affinity, hydrophobic contacts, desolvation and entropy penalization

We want to start the analysis with the ligand features that correlate with affinity, and of course we can focus first on the better and worse binders in both upper and lower ring substitutions (**Figures 2 & 3, Table S1**). Best binders in the upper ring exhibit aromatic groups with polar heteroatom substitutions, while the worse ones in this same upper-ring group exhibit, on average bulky groups or charged-ones. Interestingly, the trend remains similar in the lower ring: In relative terms, the best binders in this group are much smaller groups with sometimes, heteroatoms such fluorine, while the worse binders (and in the whole dataset) display bulky-polarizable groups, with sometimes, a fixed positive charge, such as saturated heterocyclic (nitrogen-substituted) rings.

We can analyze further the results of the changes in the water coordination number (i), the fraction of contacts in the pocket (ii), and the entropy penalization upon binding (iii) on a similar direction (**Figures 6, S9 & S10, Tables S2 & S3**): For instance, in (i), in the upper ring, derivatives that in relative terms, lose more water molecules upon binding, correspond to either bulky, or aromatic-polar-substituted ones, while those with least desolvation penalty present smaller groups or more lipophilic ones. This trend preserves in the lower ring derivatives: Bigger molecules, i.e. with more solvent-exposed surface, or polar ones, tend to have more desolvation penalty, while lipophilic ones exhibit less change in water coordination upon binding. Interestingly, the best binders lie on the region in which the penalty of losing waters is high, but not extreme, since still good binders such as 253h lie in the middle zone of relative desolvation.

Taken together, these results suggest that losing waters upon binding correlate on average with improved affinity, which in turn indicates that, the cost of removing waters around a derivative, does not hinder to a greater extent the affinity of the molecule toward the channel, likely compensated by other interactions. Still, the fact that good binders have a relative medium change in coordination number, indicates that this relationship is not in the full desolvation penalty regime, but still leaves room for some molecules that can retain lesser desolvation penalty and still exhibit good affinity.

Now, in (ii) (**Figures 6 & S9, Table S2**), focusing on the hydrophobic-hydrophilic fraction of contacts difference, one can apply a similar approach. The comparison is made in relative terms, since all derivatives exhibit more hydrophobic contact than polar ones under our definition (positive values), but still, relative differences within molecules can be made: Upper ring substitutions exhibit relatively more hydrophobic than polar contact; likewise the lower ring derivatives. Still, once again the best binders lie in the middle of this spectrum of contacts difference.

Finally, our proxy for entropy (iii) (**Figures 6 & S10, Table S3**) penalty upon binding can be also disentangled in function of substitutions: The less entropy-penalized derivatives in the upper ring substitutions display disparate trends, with bulky substituents, small and hydrophobic, or aromatic with polar heteroatoms; while the lower-ring ones show bulky groups, and planar aromatic groups with polar heteroatoms. Meanwhile, upper ring substitutions that display the greatest penalization have either short and medium-polar groups attached to the phenyl ring; or a bulkier and more polar group. In the case of the lower-ring substitutions, medium-polar groups are likewise associated with the greatest penalization.

The best binders lie more in the spectrum of high entropy penalization although, once again, not completely in the extreme values of the dataset, but around -30 J/molK, on average, compared to more extreme values such as -32-33 J/molK (**Figure S10**). Interestingly, derivatives with longer and more planar substituents on either the upper or lower ring, such as 252h and 66t, display relatively small dihedral penalties despite their poor binding affinities. This suggests that the conformations sampled along these dihedrals in solution are similar to those sampled in the bound state. Rather than indicating favorable binding, however, this behavior may reflect a relatively weak conformational restriction within the pocket: these ligands may not adopt a well-defined binding pose and may therefore retain substantial rotational freedom around the amide linkage. In general, restricting a ligand to a narrower conformational ensemble upon binding is expected to contribute unfavorably to the binding free energy through the loss of conformational entropy ($-T\Delta S$). Such a penalty can nevertheless be compensated by favorable enthalpic interactions with the protein and solvent, as well as by entropy changes associated with other ligand, protein, or solvent degrees of freedom that are not captured by the dihedrals analyzed here.
